# FBW7 couples structural integrity with functional output of primary cilia

**DOI:** 10.1101/2020.12.18.423369

**Authors:** Eleni Petsouki, Vasileios Gerakopoulos, Nicholas Szeto, Wenhan Chang, Mary Beth Humphrey, Leonidas Tsiokas

## Abstract

Structural defects in cilia have robust effects in diverse tissues and systems. However, how ciliary length changes influence signaling output are unknown. Here, we examined the functional role of a ciliary length control mechanism whereby FBW7-mediated destruction of NDE1 positively regulated ciliary length, in mesenchymal stem cell differentiation. We show that FBW7 functions as a master regulator of both negative (NDE1) and positive (TALPID3) regulators of ciliogenesis, with an overall positive net effect on cilia formation, MSC differentiation, and bone architecture. Deletion of *Fbxw7* suppresses ciliation, Hedgehog activity, and differentiation, which are rescued in *Fbxw7/Nde1*-null cells. However, despite formation of abnormally long cilia in *Nde1*-null cells, MSC differentiation is suppressed. NDE1 promotes MSC differentiation by increasing the activity of the Hedgehog pathway by direct binding and enhancing GLI2 activity in a cilia-independent manner. We propose that ciliary structure-function coupling is determined by intricate interactions of structural and functional proteins.

## Introduction

The primary cilium is a solitary, antenna-like, sensory organelle protruding into the extracellular space ^1^. It is present in virtually all cell types of the human body functioning as a signaling center for receptor tyrosine kinases, G protein coupled receptors, the Hedgehog, Notch, and Wnt pathways ^2-6^. A unique feature of the primary cilium is its ability to oscillate out of phase with the cell cycle, as primary cilia are formed when cells exit the cell cycle, whereas they start disassembling upon entry into the cell cycle ^7-12^. This oscillatory pattern of cilia enables cells to coordinate multiple signaling pathways with the cell cycle. It has been suggested that changes in ciliary length may influence the output of cilia signaling pathways ^13, 14^, but the mechanisms underlying this coordination of cilia structure and function are complex and not well understood.

One of the signaling pathways that requires intact primary cilia for maximal activity is the Hedgehog pathway ^15, 16^. In the absence of the Hedgehog ligand, GLI2 and GLI3 transcription factors are proteolytically processed at the base of the cilium to generate transcriptional repressors, GLI2R and GLI3R. In the presence of Hedgehog ligand, proteolytic cleavage is inhibited allowing the accumulation of full length GLI2 and GLI3 (GLI2A or GLI3A) that initially translocate to the tip of the cilium and subsequently in the nucleus, where they activate transcription of target genes ^5, 17^. Some target genes are activated by simple removal of GLI2R and/or GLI3R from their promoters, while others require transactivation mediated by GLI2A and/or GLI3A. Although all details of pathway activation have not been completely defined, it is well-established that the primary cilium provides the structural framework for maximal Hedgehog activity in vertebrates. This property makes it an ideal pathway to study mechanisms of coupling of ciliary structure and signaling output. Because of fundamental roles of the Hedgehog pathway in multiple systems, including the skeletal system and especially mesenchymal stem cell differentiation to osteoblasts and/or chondrocytes ^18^, functionality of this pathway can be further used in the context of stem cell differentiation in response to changes in ciliary structure.

Bone marrow derived MSCs are a progenitor cell source that has the ability to differentiate into multiple cell lineages, including osteoblasts, adipocytes, and chondrocytes ^19-21^. In addition, MSCs contribute to the formation of the niche for hematopoietic stem cells in the bone marrow, which give rise to all blood cell types. Reciprocal interactions between MSCs and HSCs govern the differentiation process of numerous cell types and defects in MSC differentiation can have widespread effects ranging from osteopenia and chondrodysplasias to obesity and cancer metastasis. MSCs possess a primary cilium and previous work has shown that it is essential for the differentiation of these cells to osteoblasts ^22^. However, exact effects of cilia *per se*, ciliary length, and contribution of cilia-based signalling and their underlying mechanisms in MSC differentiation are not completely understood.

We have identified an endogenous program regulating ciliary length in terminally differentiated cells ^13^. This program involves FBW7, the recognition receptor of the SCF^FBW7^ E3 ubiquitin ligase. FBW7 has established roles in maintenance, self-renewal and differentiation of several adult stem cell types including cancer initiating cells ^23, 24^, but the contribution of cilia to these effects is unknown. We previously showed that FBW7 mediates its effect on ciliary length by the timely destruction of NDE1, a negative regulator of ciliogenesis ^25^, upon cell cycle exit. NDE1 is phosphorylated by CDK5 at a specific site, which is in turn recognized by FBW7 ligase and targeted for degradation through the Ubiquitin-proteasomal system (UPS). The gradual destruction of NDE1 allows ciliogenesis to proceed normally and cilia to reach their appropriate length ^13^. In the present study, we wished to understand how FBW7 and its target, NDE1, can affect cilia function and whether FBW7 mediates its effects on stem cell differentiation via its effects on cilia. We find that deletion of *Fbxw7* in MSCs suppressed osteoblast differentiation. Consistently postnatal deletion of *Fbxw7* in mice resulted in reduced bone formation and mild osteopenia in 3-month-old mice. Mechanistic experiments expand the role of FBW7 to the proteasomal degradation of a positive regulator of ciliogenesis (TALPID3) in addition to NDE1. TALPID3 is present at the mother centriole, positively regulates the Hedgehog pathway, and is required for ciliogenesis ^26-29^. NDE1 is also present at the mother centriole, its effect on the Hedgehog pathway had been unknown, and is a negative regulator of ciliogenesis. We show that NDE1 physically interacts with GLI2 and increases the transcriptional activity of the Hedgehog pathway. These data suggest that mechanisms controlling ciliary length and signalling are tightly intertwined, maintaining both structural and functional integrity of primary cilia that is essential in osteoblastogenesis. Because FBW7 is one of the most commonly mutated tumour suppressors, TALPID3, a gene mutated in patients with Joubert Syndrome, and NDE1 a microcephaly gene, our data have implications in understanding the pathophysiology of all these diseases which although, they seem unrelated, they may share a common root in defective ciliary structure-function coupling.

## Results

### Postnatal deletion of *Fbxw7* leads to changes in bone architecture

The role of *Fbxw7* in osteoblast differentiation *in vivo* has not been determined. Because of embryonic lethality of *Fbxw7*-null mice ^30^, we used the tamoxifen-inducible *UbcCre*^*ERT2*^ driver to delete *Fbxw7* later, in postnatal mice and examined bone architecture of the distal femur in 12-week-old male mice using microcomputed tomography (μCT). Bone volume fraction (BV/TV), connectivity density and trabecular number (Tb.N), were significantly decreased (Fig. 1a-d), whereas trabecular separation (Tb.Sp), a parameter that reflects overall space between trabeculae, was increased (Fig. 1e). No difference between wild type and mutant mice was detected in trabecular thickness (Tb.Th) (Fig 1f). Serum bone-specific alkaline phosphatase levels (BALP) (Fig. 1g), but not Tartrate resistant acid phosphatase (TRAcP) (Fig. 1h) were reduced in mutant mice indicating that reduced bone mass may have been primarily caused by reduced osteoblast differentiation and/or function.

**Figure 1.**
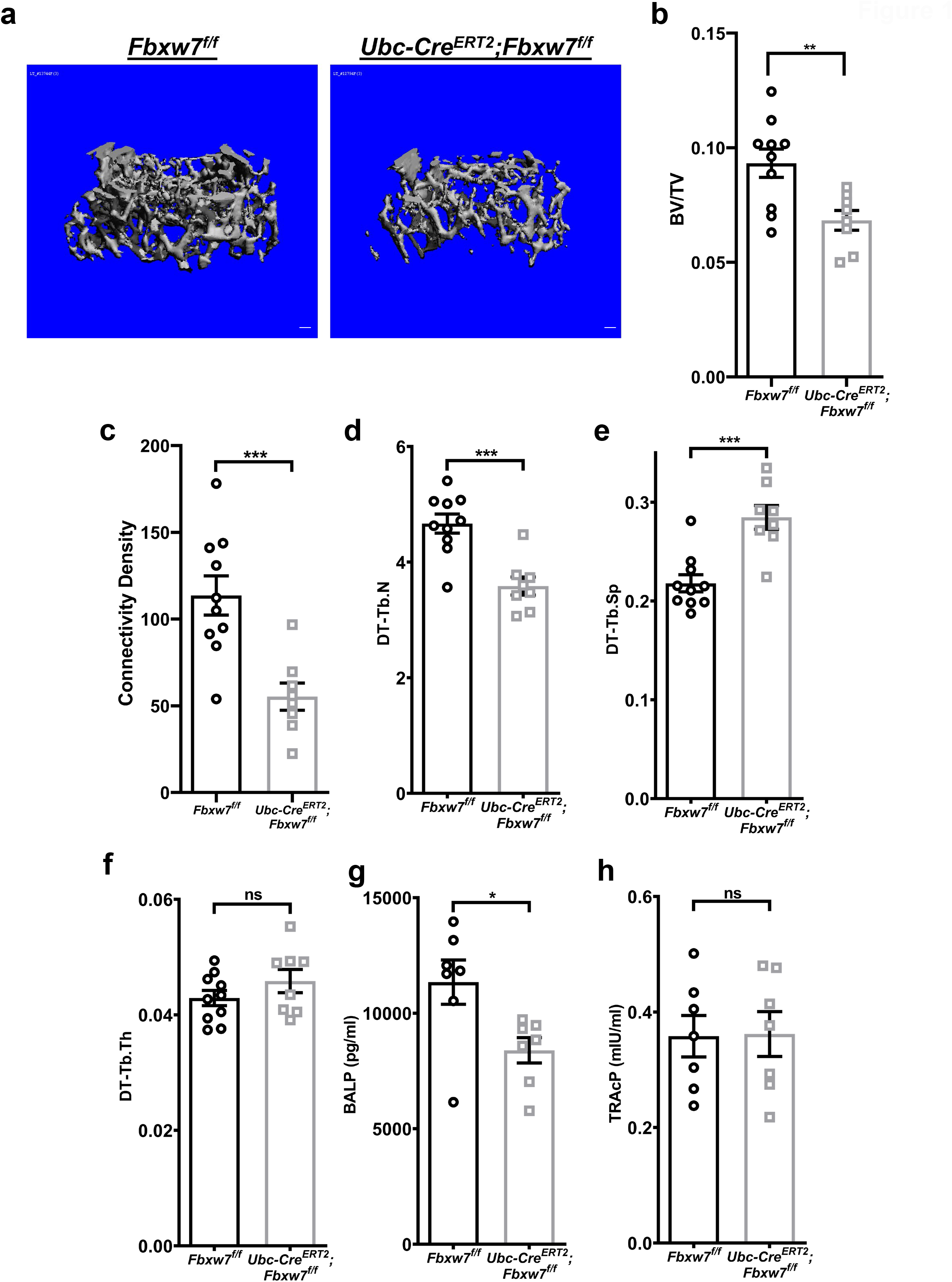
Postnatal deletion of *Fbxw7* leads to changes in bone architecture. **a)** Representative three-dimensional reconstitution analysis of the architecture of metaphyseal trabecular bone in the distal femur of 12-week-old *Fbxw7*^*f/f*^ or *UbcCreERT2; Fbxw7*^*f/f*^ mice treated with 4-Hydroxy-tamoxifen (4OHT). Scale bars: 100 µm. **b-f)** Quantitative μCT analysis of skeletal parameters: BV/TV (Bone volume fraction) (b), Connectivity density (c), DT-Tb.N (Trabecular number) (d), DT-Tb.Sp (Trabecular separation) (e) and DT-Tb.Th (Trabecular thickness) (f) of the metaphyseal trabecular bone in the distal femur of 12-week-old *Fbxw7*^*f/f*^ (n=10) or *UbcCre*^*ERT2*^;*Fbxw7*^*f/f*^ (n=8) mice treated with 4OHT. Data are presented as means ± SEM. Student’s t-test, **p < 0.01, ***p < 0.001. **g-h)** ELISA analysis for bone formation (Mouse Bone-specific alkaline phosphatase, BALP) (g) or bone resorption (Mouse Tartrate-resistant acid phosphatase 5b, TRAcP-5b) (h) markers in the serum of 12-week-old *Fbxw7*^*f/f*^ (n=7) or *UbcCreERT2; Fbxw7*^*f/f*^ (n=7) mice treated with 4OHT. Data are presented as means ± SEM. Student’s t-test, *p < 0.05.

### Deletion of *Fbxw7* reduces ciliation in MSCs or MSC-like cell types

*Ex vivo* cultures of MSCs derived from the bone marrow of *UbcCre*^*ERT2*^;*Fbxw7*^*f/f*^ mice were used to test whether deletion of *Fbxw7* can affect ciliation and differentiation of MSCs to osteoblasts. In terms of ciliation, we specifically examined CD106+ MSCs, called skeletal stem cells (SSCs), which give rise to osteoblasts, adipocytes, and chondrocytes ^31, 32 33, 34^. *Ex vivo* deletion of *Fbxw7* via 4-hyrdroxytamoxifen (4-OHT) treatment in serum starved primary cultures of mouse SSCs (CD106^+^ MSCs) resulted in the reduction of the percentage of ciliated cells compared to untreated cells (Fig. 2a,b). Because MSC cultures can be highly heterogeneous, we also examined the effect of FBW7 depletion using RNAi in transiently transfected C3H10T1/2 cells, an MSC-like stem cell line ^35, 36^. Depletion of FBW7 led not only to a reduction in the percentage of ciliated cells, but also a reduction in ciliary length in remaining ciliated cells (Fig. 2e-g).

**Figure 2.**
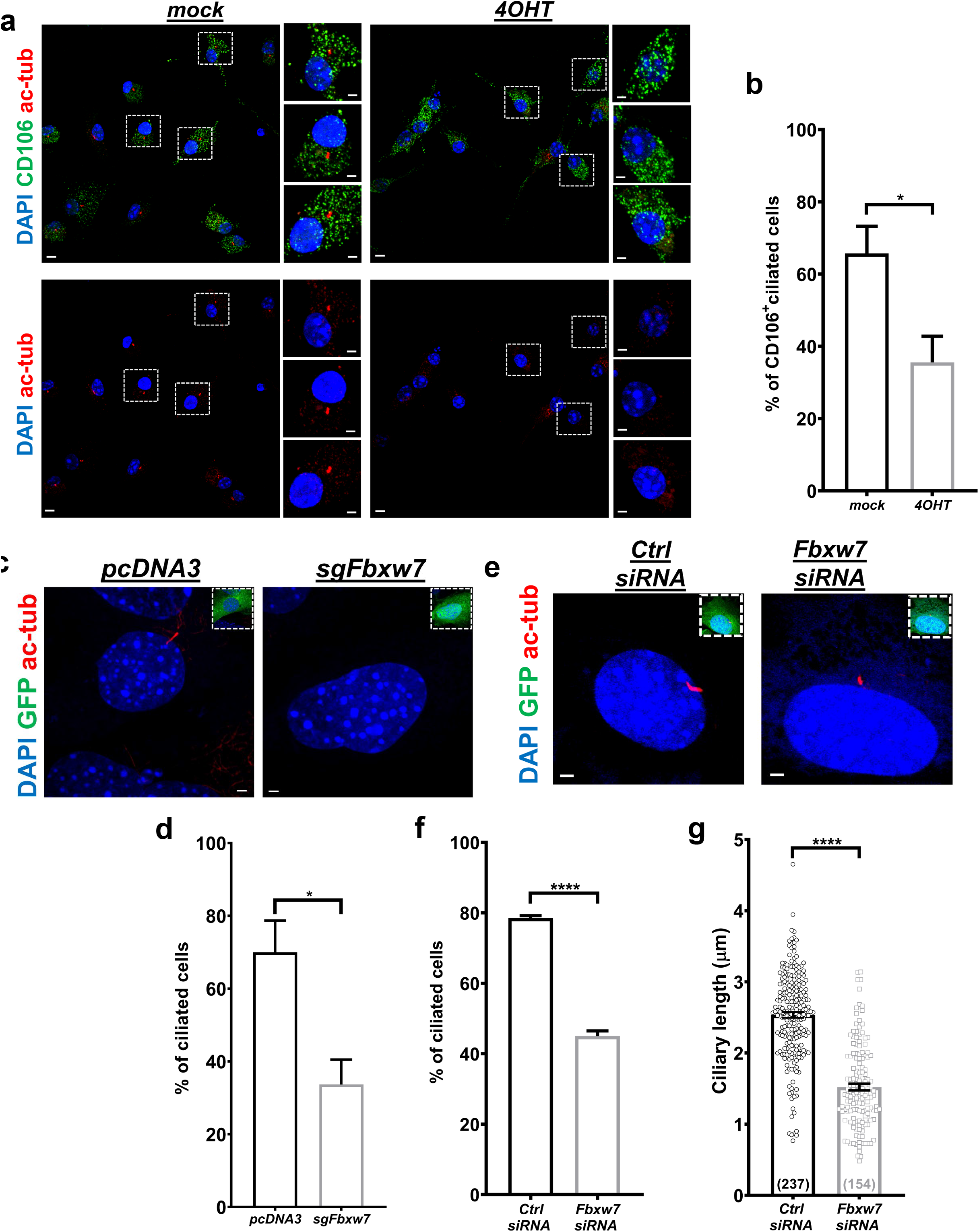
Deletion of *Fbxw7* reduces ciliation in MSCs, MEFs and C3H10T1/2 cells. **a-b)** Representative images of MSCs isolated from the bone marrow of *UbcCre*^*ERT2*^; *Fbxw7*^*f/f*^ mice, *ex vivo* treated with ethanol (mock) or 4OHT in ethanol, serum starved for 24h and stained for the SSC marker CD106 (green) and primary cilia (acetylated α-tubulin, red) (a). Percent of CD106+ cells with cilia in mock or 4OHT treated cells isolated from 4 different *UbcCre*^*ERT2*^; *Fbxw7*^*f/f*^ mice at 4 weeks of age (b). 100 mock- and 100 4OHT-treated CD106^+^ SSCs were analyzed per mouse. Scale bars: 5 µm. Scale bars in insets: 2 µm. Data are presented as means ± SEM. Student’s t-test, *p < 0.05. **c-d)** Representative images of mouse embryonic fibroblasts (MEFs) transfected with GFP and pcDNA3 or *Fbxw7*-specific sgRNA (sg*Fbxw7*), serum starved for 24h and stained for primary cilia (acetylated α-tubulin, red) (c). Percent of ciliated MEFs treated with the indicated constructs **d)**. 100 GFP^+^ MEFs were analyzed for each group per experiment (n=3). Scale bars: 2 µm. Data are presented as means ± SEM. Student’s t-test, *p < 0.05. **e)** Representative images of C3H10T1/2 cells transfected with GFP and the indicated constructs, and serum starved for 24h. Scale bars: 2 µm. **f**,**g)** Percent of ciliated cells (f, n=3) and ciliary length (g) of cells in (e). Data are presented as means ± SEM. Student’s t-test, ****p < 0.0001.

Reduced levels of ciliation were also observed in mouse embryonic fibroblasts (MEFS) transiently transfected with a *Fbxw7*-specific sgRNA construct (Fig. 2c,d).These data were consistent with our previous results in hTERT-RPE1 cells ^13^, that FBW7 has an essential role in ciliary length control.

### *Fbxw7* deletion suppresses differentiation of MSCs to osteoblasts

Primary cultures of *UbcCre*^*ERT2*^; *Fbxw7*^*fl/fl*^ MSCs allowed us to delete *Fbxw7* at either a stem cell stage by inducing deletion before treatment with differentiation media (Fig. 3a), or at a stage where cells had already been committed to differentiate, by inducing deletion after addition of differentiation media (Supplementary Fig. 1). Deletion of *Fbxw7* in MSCs before induction of differentiation resulted in significant suppression of osteoblast differentiation (Fig. 3b), as demonstrated by Alizarin red S staining. In addition, mRNA levels of four out of five osteogenic markers examined were significantly reduced at all time-points after osteogenic induction (Fig. 3c-e). On the other hand, deletion of *Fbxw7* after osteogenic induction resulted in a milder suppression of differentiation (Supplementary Fig. 1). Tamoxifen treatment did not have an effect on cells in which *Fbxw7* was not floxed (Supplementary Fig. 2). In sum, these results suggested that FBW7 promoted osteoblast differentiation by acting predominantly at the stage of osteoblast lineage commitment rather than by supporting maintenance of already committed pre-osteoblasts.

**Figure 3.**
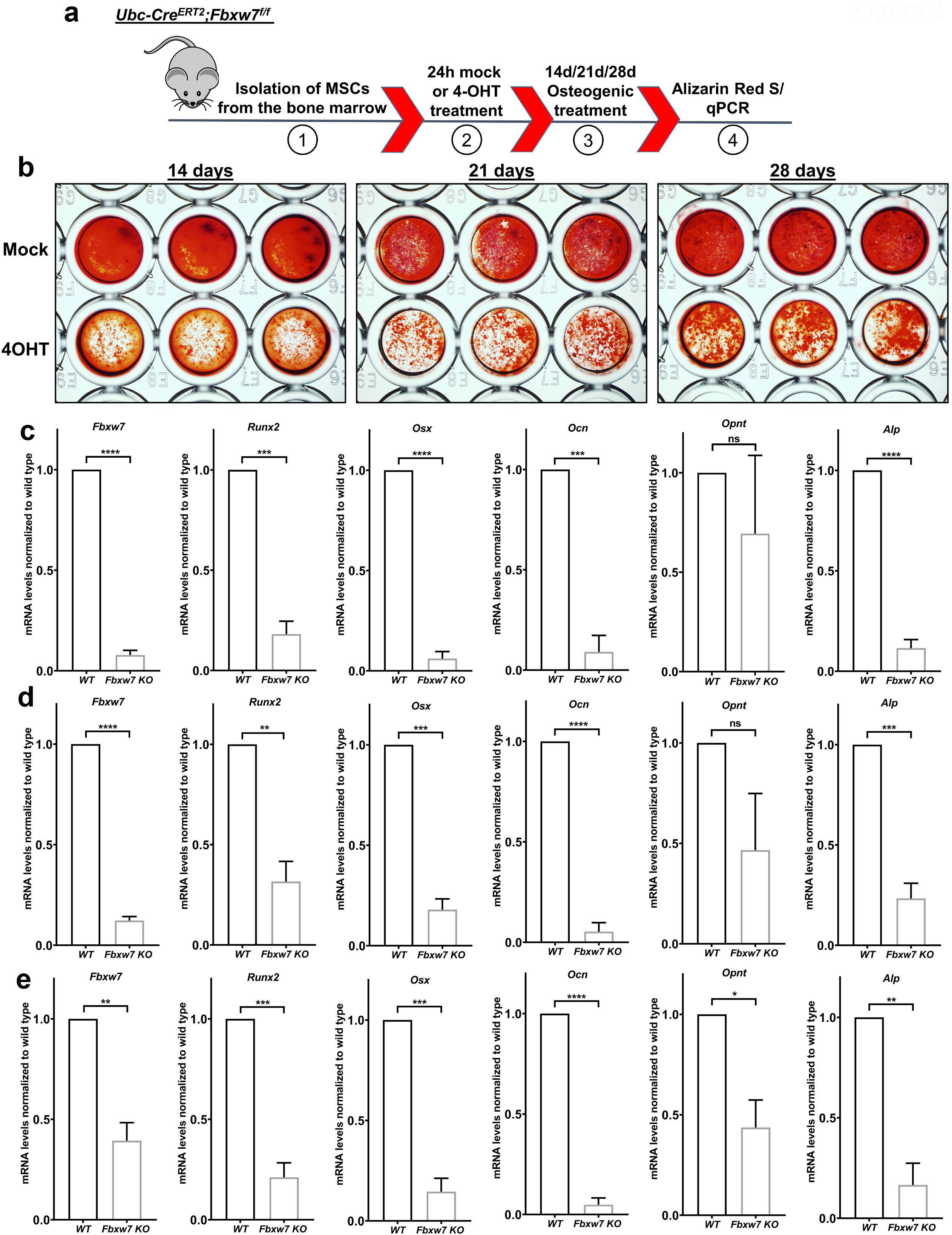
*Fbxw7* deletion before osteogenic induction severely suppresses differentiation of MSCs to osteoblasts. **a)** Diagram showing an experimental setup for *ex vivo* deletion of *Fbxw7*. Isolation of MSCs from the bone marrow of *UbcCre*^*ERT2*^; *Fbxw7*^*f/f*^ mice (1) was followed by 4OHT-induced deletion of *Fbxw7* (2), osteogenic induction (3) and analysis of osteoblast differentiation (4). **b-e)** Osteoblast differentiation of *UbcCre*^*ERT2*^; *Fbxw7*^*f/f*^-derived MSCs treated with mock or 4OHT before osteogenic induction (n=3 different mice). Differentiation was measured at 14, 21 and 28 days after osteogenic induction via Alizarin Red S staining (b) and mRNA levels of osteoblast differentiation markers *Runx2, Osterix* (*Osx*), *Osteocalcin* (Ocn), *Osteopontin* (*Opnt*) and Alkaline phosphatase (*Alp*) at 14 (c), 21 (d) and 28 days (e) after osteogenic induction. mRNA levels of *Fbxw7* were analyzed to confirm deletion of the gene. Data are presented as means ± SEM. Student’s t-test, *p < 0.05, **p < 0.01, ***p < 0.001, ****p < 0.0001.

In contrast to what was seen in osteoblasts, deletion *of Fbxw7* before adipogenic treatment promoted adipogenic differentiation of MSCs isolated from *UbcCre*^*ERT2*^; *Fbxw7*^*fl/fl*^ mice (Supplementary Fig. 3). However, deletion of *Fbxw7* after commitment to adipogenesis showed a trend to increase differentiation without reaching statistical significance (Supplementary Fig. 3). These results indicated that deletion of *Fbxw7* did not generally impair the ability of MSCs to differentiate, but it rather skewed differentiation towards the adipogenic lineage at the expense of osteoblastic differentiation. This type of effect suggested that FBW7 contributes to stem cell lineage determination rather than affecting committed cell types.

### FBW7 regulates ciliary length through NDE1 in the C3H10T1/2 stem cell line

Since we established a cellular role of FBW7 in osteoblast differentiation, we next determined whether this role was mediated via an effect on cilia and NDE1. Thus, first we tested whether deletion of both *Nde1* and *Fbxw7* would rescue reduced ciliation and ciliary length induced by loss of *Fbxw7* alone, as previously shown in hTERT-RPE1 cells ^13^. This hypothesis was tested in the cell line C3H10T1/2 for several reasons. First, these cells are multipotent and have been extensively used as a surrogate cell culture system to model osteoblast, adipocyte, or chondrocyte differentiation in vitro ^35, 36^. Second, they express FBW7 and NDE1, but not NDEL1 (Supplementary Fig. 4). NDEL1 is a homolog of NDE1 that functions redundantly to NDE1 in ciliogenesis ^37, 38^. This effect has been described *in vivo* and is believed to account for the much milder phenotype of *Nde1*-null compared to *Ndel1*-null mice restricted only to neuronal cell types ^39^. Most relevant to our study, NDEL1 is not a substrate of FBW7 and therefore it would evade FBW7-mediated degradation. Finally, the Hedgehog pathway drives osteoblast differentiation of C3H10T1/2 cells and therefore, cilia-based signaling is functionally relevant in these cells ^40, 41^.

**Figure 4.**
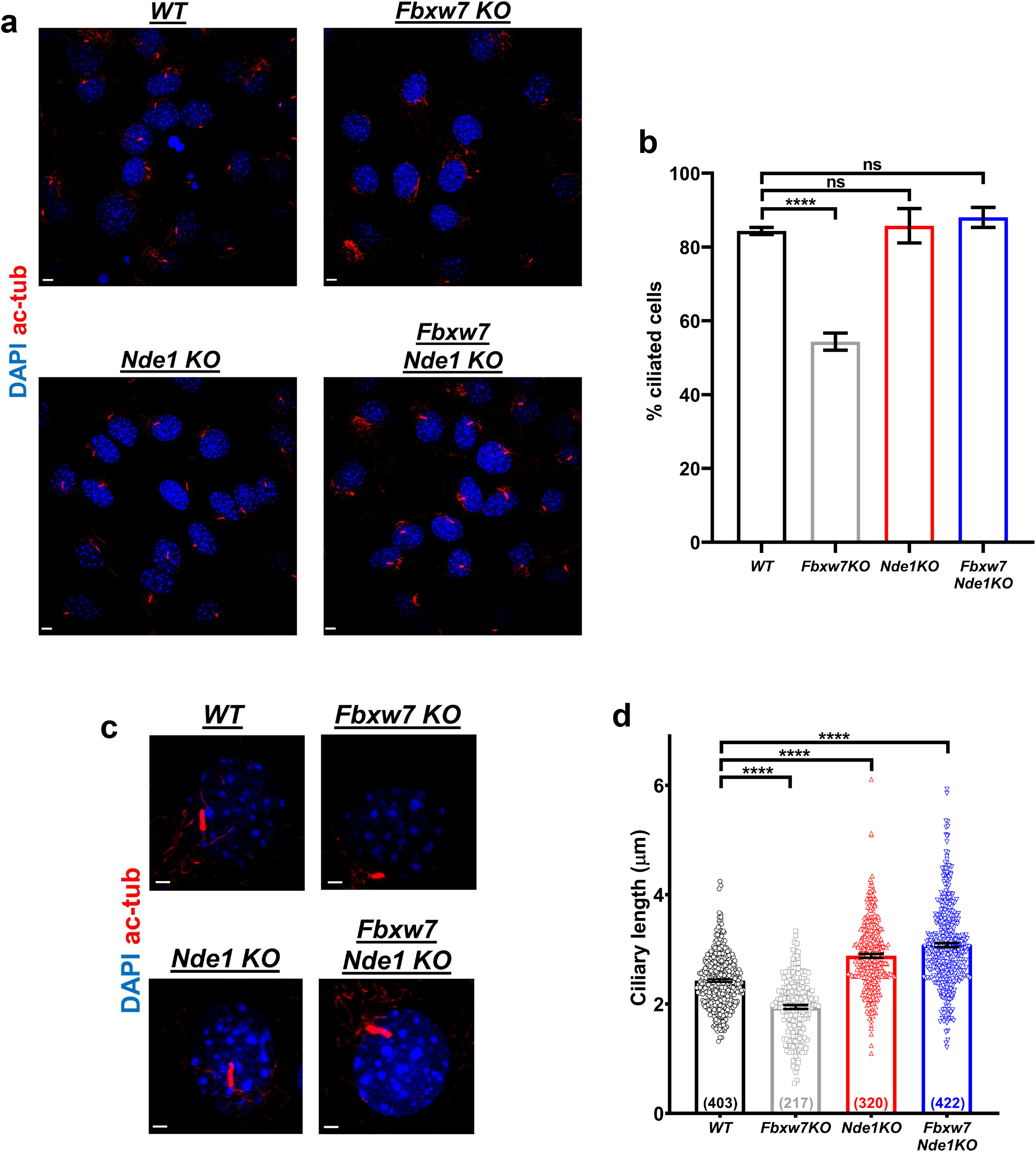
FBW7 regulates ciliation and ciliary length through NDE1 in the C3H10T1/2 stem cell line. **a-b)** Ciliation analysis of wild type, *Fbxw7KO, Nde1KO* and *Fbxw7Nde1KO* C3H10T1/2 cells after 24h of serum starvation. Representative images of the indicated cells stained for primary cilia (acetylated α-tubulin, red) (a). Percent of ciliation in the indicated C3H10T1/2 cells (b). 100-150 cells were analyzed per experiment (n=3). Scale bars: 5 µm. Data are presented as means ± SEM. One-way ANOVA with Dunnett’s multiple comparisons test, ****p < 0.0001. **c-d)** Ciliary length analysis of wild type, *Fbxw7KO, Nde1KO* and *Fbxw7Nde1KO* C3H10T1/2 cells after 24h of serum starvation. Representative images of the indicated cells stained for primary cilia (acetylated α-tubulin, red) (c). Ciliary length in the indicated C3H10T1/2 cells (d). The number of cells measured per group is indicated at the bottom of each bar. Scale bars: 2 µm. Data are presented as means ± SEM. One-way ANOVA with Dunnett’s multiple comparisons test, ****p < 0.0001.

We generated three stable lines, lacking *Fbxw7, Nde1*, or both using CRISPR/Cas9 gene editing. Inactivation of these genes was confirmed by both Sanger sequencing of the edited locus and protein analysis (Supplementary Fig. 4). Because current batches of the NDE1 antibody recognize an additional band in mouse lysates, we used MEFs from *Nde1*^*-/-*^ mice to identify the band corresponded to NDE1. The lower band detected by the NDE1 antibody corresponded to NDE1 (Supplementary Fig. 4). Cells lacking *Fbxw7* showed a reduction in the percentage of ciliated cells and ciliary length in remaining ciliated cells (Fig. 4), as shown in SSCs (Fig. 2a,b). Deletion of *Nde1* did not increase the percentage of ciliated cells 24 h after serum starvation, but rather ciliary length, as we had previously shown in NIH3T3 and hTERT-RPE1 cells ^25^. In double mutant cells, the percentage of ciliated cells was similar to wild type levels, while ciliary length was significantly higher than wild type cells.

### FBW7 regulates functional integrity of cilia in C3H10T1/2 cells

While experiments above showed that the FBW7/NDE1 module is essential for cilia structure, they could not inform us about cilia function. Thus, we tested whether cilia *per se* are essential for osteoblast differentiation. We inhibited cilia formation by two means, CRISPR/Cas9-mediated inactivation of the *Ift88* gene or treatment of cells with Ciliobrevin A, an inhibitor of cytoplasmic dynein, Hedgehog pathway, and cilia formation ^42^. Cells lacking cilia failed to differentiate to osteoblast-like cells, as determined by expression of alkaline phosphatase (ALP), a well-established marker of osteoblast differentiation in these and other multipotent cells (Supplementary Fig. 5). Osteoblast differentiation was severely reduced in single *Fbxw7* and *Nde1*-deleted cells compared to wild type cells, but partially restored in double mutant cells (Fig. 5a-d). In addition, we used MLN4924, an inhibitor of Cullin-dependent ligases, including FBW7 ^43^. Initiation of treatment of wild type C3H10T1/2 cells with MLN4924 before osteogenic induction resulted in suppression of differentiation. However, initiation of MLN4924 treatment only after osteogenic induction had no effect on differentiation (Supplementary Fig. 6). These data were consistent with data using 4-OHT-inducible deletion of *Fbxw7* in MSCs (Fig. 3 & Supplementary Fig. 1), suggesting that SCF^FBW7^ activity at the pre-commitment stage, but not after commitment, is more relevant to osteoblast differentiation. Further, because effects of single knockouts were partially corrected in double mutant cells, these results supported the hypothesis that the FBW7-mediated degradation of NDE1 is essential for osteoblast differentiation. To test whether cilia were functional in double mutant cells, cells were treated with Ciliobrevin A (Fig. 5f-h) or transiently transfected with an *Ift88*-specific sgRNA (Fig. 5e). Indeed, ALP levels were dramatically suppressed in these cells, suggesting that deletion of *Fbxw7* and *Nde1* led to the formation of functional cilia (Fig. 5). While the reduced level of differentiation in FBW7 depleted cells could be consistent with the reduced percentage of ciliated cells and ciliary length, the reduced level of differentiation in *Nde1*-null cells was unexpected based on effects on cilia alone and suggested that knocking out NDE1 uncoupled structure from function. Similar results were obtained in another clone of C3H10T1/2 cells lacking *Nde1*, ruling out clonal effects. Based on these data, we reasoned that while deletion of *Nde1* led to abnormally long cilia, these cilia were not functional, yet they acquire some level of functionality in double mutant cells.

**Figure 5.**
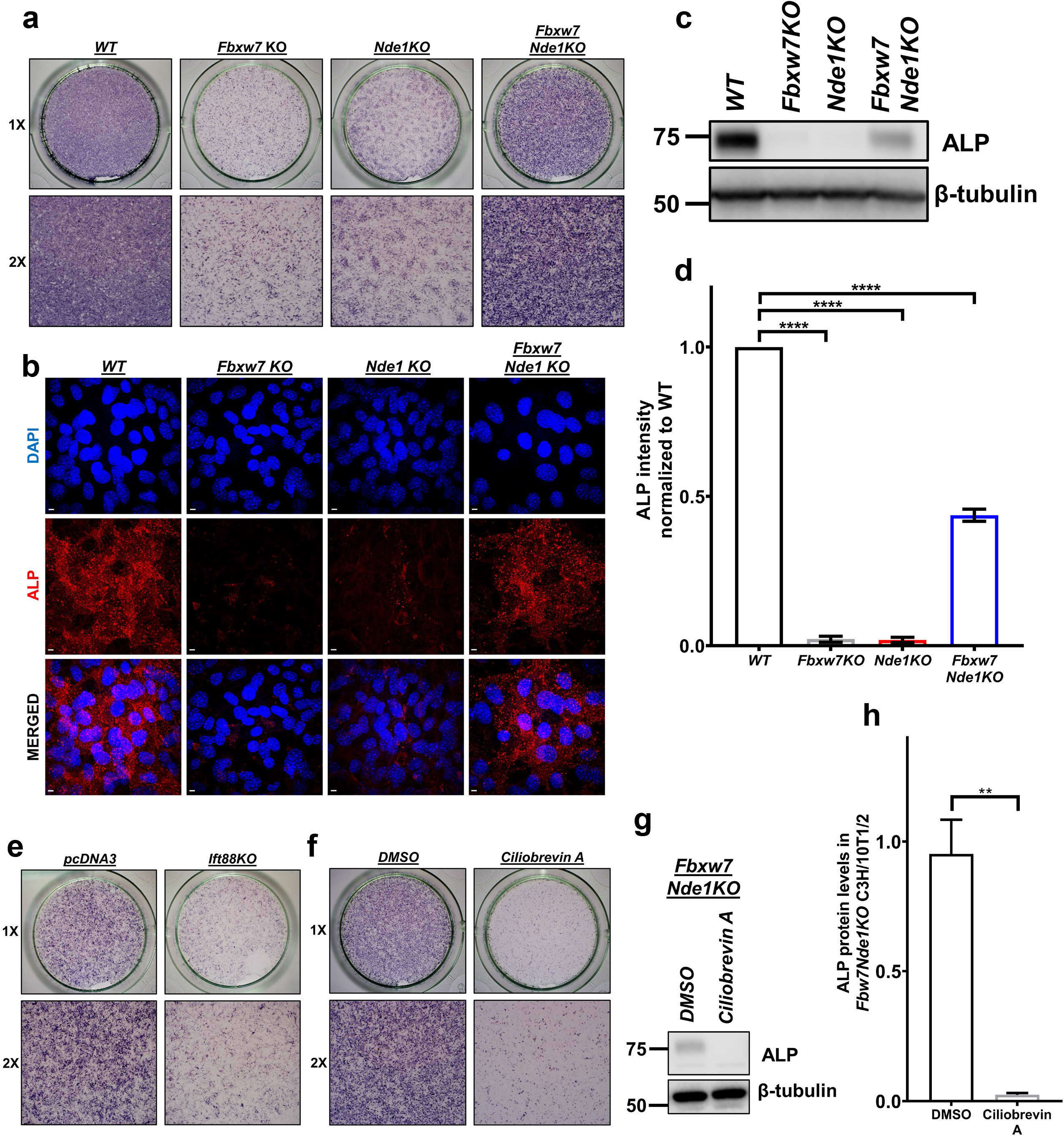
FBW7-NDE1 axis affects osteoblast differentiation via effects on primary cilia in C3H10T1/2 cells. **a**,**b)** Representative images for ALP staining (a) and ALP immunofluorescence (red) (b) of wild type, *Fbxw7KO, Nde1KO* and *Fbxw7Nde1KO* C3H10T1/2 cells (n=3). 2X indicates magnification of corresponding 1X images. Scale bars: 5 µm. **c**,**d)** Expression levels of ALP in indicated cells (c) and summary data (d). Data are presented as means ± SEM. One-way ANOVA with Dunnett’s multiple comparisons test, ****p < 0.0001. **e**,**f)** ALP staining after transfection of *Fbxw7Nde1KO* C3H10T1/2 cells with pcDNA3 or an *Ift88*-specific sgRNA (e) or after treatment with DMSO or Ciliobrevin A (f). 2X indicates magnification of corresponding 1X images. **g**,**h)** Expression levels of ALP in *Fbxw7Nde1KO* C3H10T1/2 cells after treatment with DMSO or Ciliobrevin A (g). Summary data of (g) are shown in (h). Data are presented as means ± SEM. Student’s t-test, **p < 0.01

Considering the central role of the Hedgehog pathway in osteoblast differentiation and the dependence of this pathway on structural integrity of primary cilia ^5, 17, 18^, we transfected all four cell lines with a Gli-reporter construct and determined basal level activity of the Hedgehog pathway. As shown in Fig. 6a, pathway activity correlated with differentiation levels. We reasoned that low activity of Hedgehog in *Fbxw7-*null cells was due to decreased levels of ciliation and reduced ciliary length. However, since *Nde1*-null cells have normal ciliation and even longer primary cilia compared to wild type, we hypothesized that functionality of cilia-related Hedgehog effectors might have been compromised. Thus, we tested whether overexpression of various forms of GL12 or GLI3 could rescue differentiation in *Nde1*-null cells (Fig. 6b,c and Supplementary Fig. 5). Full length or constitutively active GLI2 increased differentiation in these cells, suggesting the possible defects in GLI2-mediated signaling may account for the impaired cilia functionality in *Nde1*-null cells ^17, 44^. Interestingly, overexpression of a constitutively active full-length GLI3 construct (GLI3 P1-P6) had no effect in *Nde1*-null cells ^17^, while acting as a dominant negative allele in *Fbxw7*-null cells (Supplementary Fig. 5), highlighting the importance and specificity of GLI2 in cilia-mediated osteoblastogenesis in this system. Here, it is interesting to note that it was not GLI3R that could account for the dominant negative effect of GLI3 P1-P6 on *Fbxw7*-null cells, because this GLI3 form is resistant to proteolytic cleavage.

**Figure 6.**
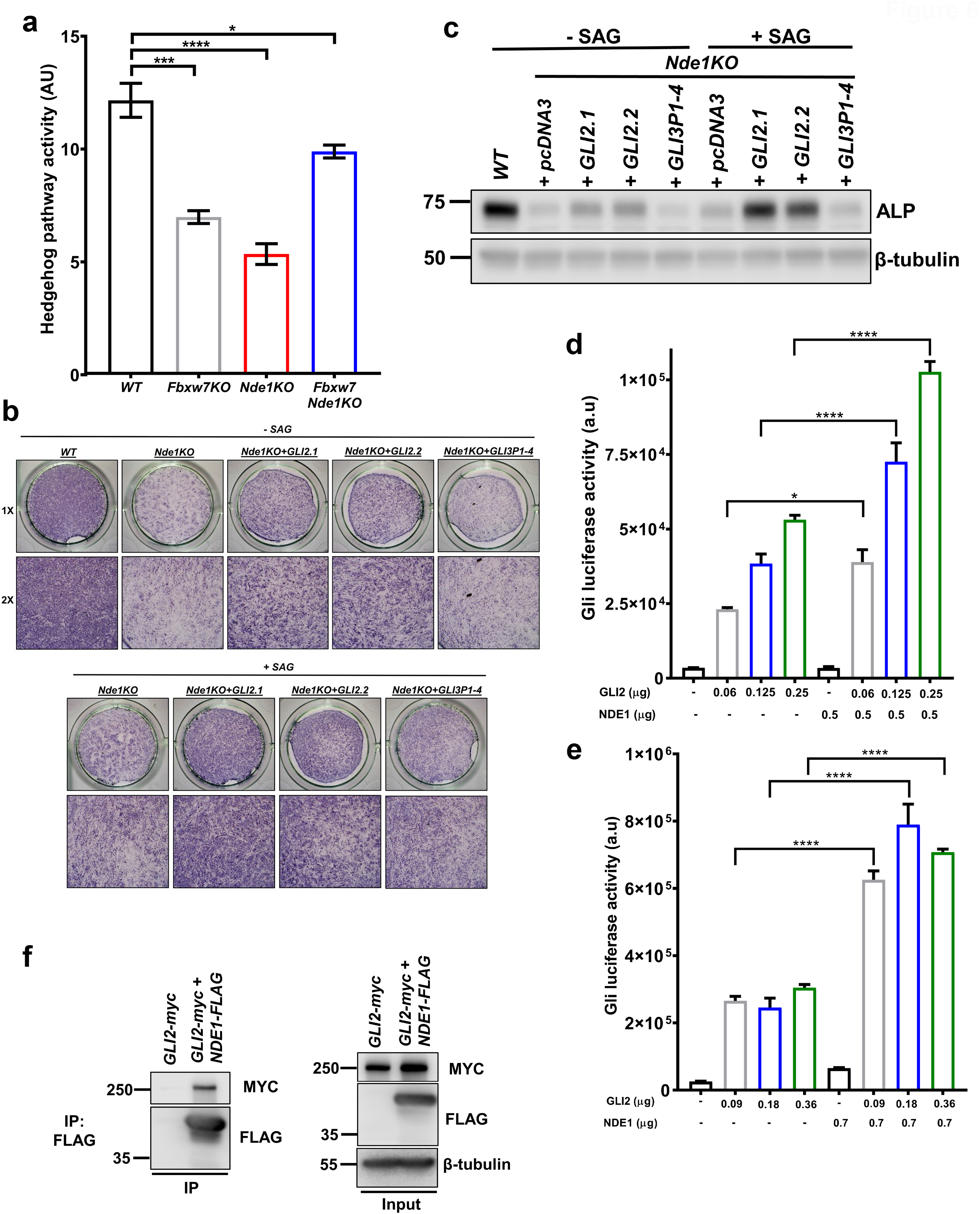
Hedgehog pathway activity in single and double mutant cells and its modulation by NDE1. **a)** Summary data of Hedgehog pathway activity measured as luciferase activity after transfection of wild type, *Fbxw7KO, Nde1KO* and *Fbxw7Nde1KO* C3H10T1/2 cells with a Gli-reporter construct (n=3). Data are presented as means ± SEM. One-way ANOVA with Dunnett’s multiple comparisons test, *p < 0.05, ***p < 0.001, ****p < 0.0001. **b**,**c)** Representative ALP staining (b) and protein expression levels (c) of wild type and *Nde1KO* C3H10T1/2 cells transfected with the indicated constructs and treated with or without Smoothened agonist (SAG). **d**,**e)** Summary data of Hedgehog pathway activity measured as luciferase activity after transfection of C3H10T1/2 (d) or HEK293T cells (e) with a Gli-reporter construct and the indicated constructs (n=3). Data are presented as means ± SEM. One-way ANOVA with Dunnett’s multiple comparisons test, ****p < 0.0001. **f)** Physical interaction of GLI2-myc and NDE1-FLAG constructs in HEK293T cells.

### NDE1 physically interacts with and increases the transcriptional activity of GLI2

Hedgehog activity was decreased in *Nde1*-null cells despite the presence of cilia. This prompted us to test whether NDE1 increased the activity of GLI2 and physically interacted with GLI2. Transient transfection in C3H10T1/2 and HEK293T cells indicated that GLI2 activity was increased in the presence of NDE1 (Fig. 6d,e) and NDE1 co-immunoprecipitated with GLI2 (Fig. 6f). These data could help explain the reduced GLI activity in *Nde1*-null cells and further indicate that while NDE1 functions as a negative regulator of ciliogenesis, it functions as a positive regulator of cilia based signaling such as Hedgehog signaling, coupling ciliary structure and function.

### Identification of TALPID3 as a possible target of FBW7

Because osteoblast differentiation was partially rescued in double mutant cells, while cilia appeared grossly structurally similar to cilia of *Nde1*-null cells, we reasoned that positive regulator(s) of the Hedgehog pathway might be additional potential targets of FBW7 that could accumulate in double mutant cells providing some degree of rescue. Screening all known positive regulators of the Hedgehog pathway for the presence of optimal FBW7 phosphodegrons, we identified Fused and TALIPD3 as potential targets. We proceeded with TALPID3, because it had 4 optimal FBW7 phosphodegrons (Fig. 7d). In addition, TALPID3 is required for early stages of cilia formation and organization of transition fibers to assemble the ciliary gate at the base of the cilium ^27^, which would be consistent with the small but significant increase in ciliary length in double mutant cells. As predicted, all FBW7 isoforms co-immunoprecipitated with TALPID3, albeit at different levels (Fig. 7a,b). Ubiquitinylation experiments in HEK293T cells suggested that co-expression of TALPID3 with wild type, but not ubiquitin ligase dead FBW7 mutants, resulted in increased ubiquitinylation of TALPID3 (Fig. 7c). Interestingly, an N-terminally truncated form of FBW7 that has been widely used for functional studies was stabilized when co-transfected with TALPID3, suggesting that binding of TALPID3 to FBW7 may have blocked its auto-ubiquitylation ^45^ supporting the idea that TALPID3 is a genuine target of FBW7 (Fig. 7a). Consistent with the positive effect of TALPID3 on Hedgehog signaling and osteoblast differentiation, transient depletion of *Talpid3* mRNA in wild type or double mutant *Fbxw7/Nde1* cells, resulted in marked reductions of both functional read-outs (Fig. 7e-l). Efficient depletion of mouse TALIPD3 by *Talpid3* siRNA was confirmed in transfected C3H10T1/3 cells (Supplementary Fig. 4). These data suggest FBW7 controls the abundance of TALPID3 in addition to NDE1 and the coordinated effect of these, and possibly other yet unidentified proteins can contribute to the cilia structure-function coupling in cells lacking *Fbxw7*.

**Figure 7.**
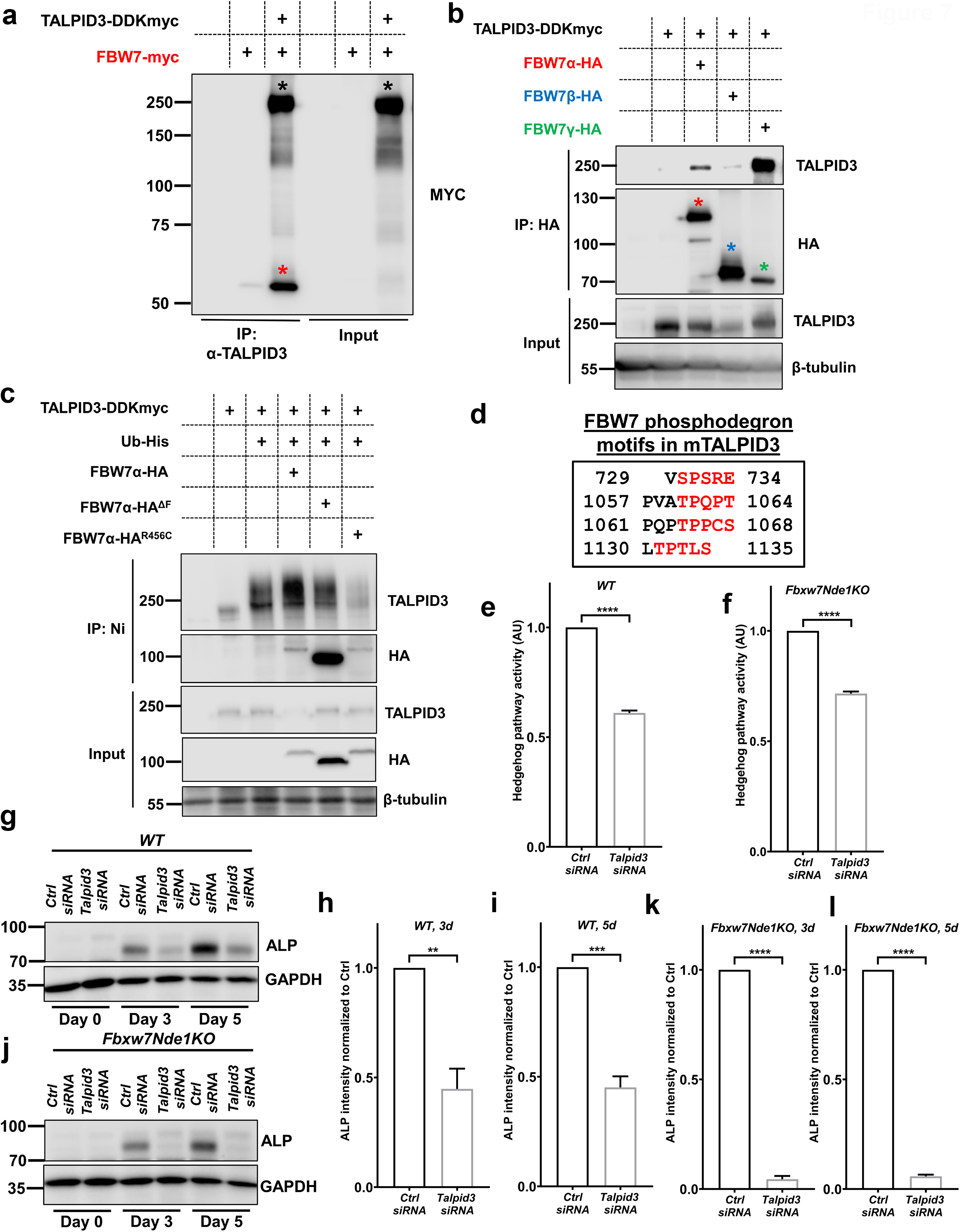
Identification of TALPID3 as a target of FBW7 and GLI2. **a**,**b)** Physical interaction of TALPID3-myc with the indicated constructs of FBW7 in HEK293T cells. Black and red asterisks in (a) indicate TALPID3-myc and FBW7-myc, respectively. Red, blue and green asterisks in (b) indicate FBW7α, -β and -γ, respectively. **c)** Ubiquitinylation assay of TALPID3-myc in HEK293T cells after co-transfection with 6-His-tagged Ubiquitin and the indicated HA-tagged FBW7 constructs. **d)** Predicted FBW7 phosphodegron motifs in mouse TALPID3. **e**,**f)** Summary data of Hedgehog pathway activity measured as luciferase activity after transfection of wild type (e) and *Fbxw7Nde1KO* (f) C3H10T1/2 cells with a Gli-reporter construct and *Talpid3* siRNA or control siRNA (n=3). Data are presented as means ± SEM. Student’s t-test, **p < 0.01, ****p < 0.0001. **g-i)** Expression levels of ALP in wild type C3H10T1/2 cells, transfected with control siRNA or *Talpid3* siRNA, at 0 days, 3 days and 5days of osteogenic differentiation (g) and summary data (h,i). Data are presented as means ± SEM. Student’s t-test, **p < 0.01, ***p < 0.001. **j-l)** Expression levels of ALP in *Fbxw7Nde1KO* C3H10T1/2 cells, transfected with control siRNA or *Talpid3* siRNA, at 0 days, 3 days and 5 days of osteogenic differentiation (j) and summary data (k,l). Data are presented as means ± SEM. Student’s t-test, ****p < 0.0001.

## Discussion

While it is known that reduction in ciliary length can result in severe phenotypes in diverse tissues and systems, it is not clear how changes in ciliary length can influence signaling output resulting in these robust phenotypes. We have shown earlier that the FBW7-mediated proteasomal degradation of NDE1 functions as a rheostat for ciliary length ^13^. Results from our present study suggest that ciliary signaling output is tuned to ciliary length by direct physical interactions of proteins that mediate structural roles with proteins mediating functional roles (Fig. 9). This conclusion is based on the following lines of evidence. First, FBW7 controls the abundance of both positive and negative regulators of ciliogenesis, TALPID3 and NDE1, respectively. Second, NDE1 physically interacts with GLI2 increasing its transcriptional activity. Third, the FBW7/NDE1/TALPID3/GLI2 network of protein-protein interactions and activities is essential for both cilia formation and differentiation of MSCs to osteoblasts and postnatal deletion of *Fbxw7* leads to reduced bone mass and osteoblast activity (Fig. 8). While the differentiation of MSCs to osteoblast was utilized as a model system to decipher molecular and cellular mechanisms of cilia structure and function, our results have implications in the pathophysiology of not only *bona-fide* ciliopathies, such as Joubert syndrome (TALPID3) ^46^, but also of emerging ciliopathies such as cancer (FBW7) ^47, 48^ and microcephaly (NDE1) ^38^.

**Figure 8.**
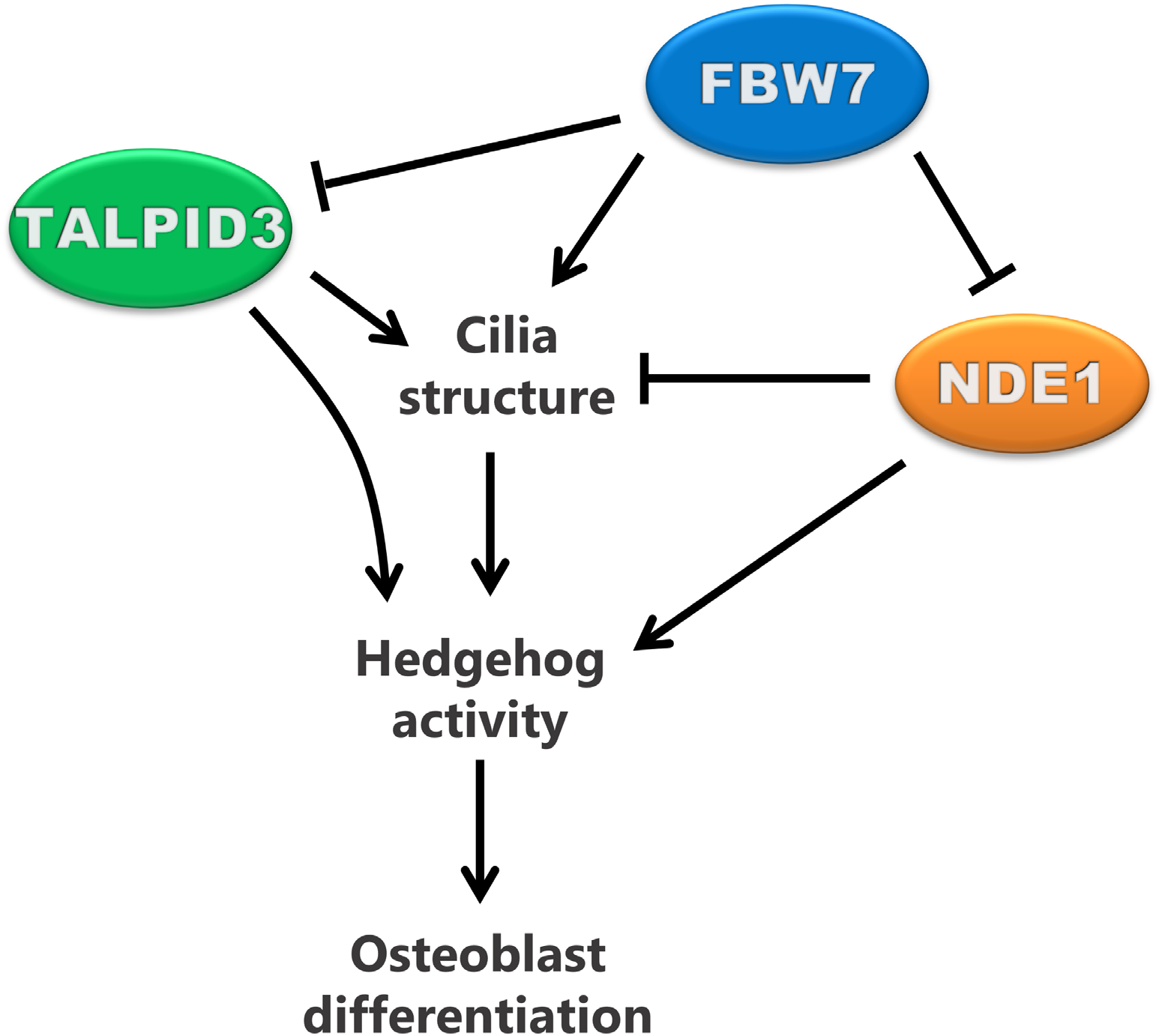
Working model. Working model depicting the role of the FBW7-NDE1-TALPID3 pathway in primary cilia structure, Hedgehog pathway activity, and osteoblast differentiation.

Our approach to mechanistically determine structure-function relationships of primary cilia entailed the identification of a functional assay for the primary cilium in cell culture that would also bear physiological relevance in the whole organism. Several lines of evidence have suggested that primary cilia are essential for the differentiation of MSCs to osteoblasts ^49-53^. Therefore, we determined whether the FBW7-mediated degradation of NDE1 can also affect ciliary function in this context. Our data showed that deletion of *Fbxw7* in primary MSCs and C3H10T1/2 suppressed both ciliation and differentiation. Consistently, postnatal deletion of *Fbxw7* in mice reduced bone mass and levels of BALP, a marker of osteoblastic activity in the serum. Double deletion of *Fbxw7* and *Nde1* in C3H10T1/2 cells rescued cilia formation and osteoblast differentiation. Treatment of the double mutants with ciliobrevin A or triple deletion of *Fbxw7, Nde1*, and *Ift88* eliminated rescued cilia and suppressed differentiation. These data lead us to propose that the FBW7/NDE1 module had a functional role in the differentiation of MSCs to osteoblasts, and such a role was mediated at least in part, via an effect on cilia. However, we unexpectedly found that cells lacking *Nde1* showed a block in differentiation despite the fact that cilia were present in these cells. These results indicated that restoring ciliary length does not automatically mean that cilia function is also restored. Further mechanistic experiments led to two key observations: 1) TALPID3, a positive regulator of ciliogenesis and Hedgehog activity, is a target of FBW7 and 2) NDE1 is a positive regulator of GLI2 and a negative regulator of ciliary length. These data lead us to propose a model in which, FBW7 controls structural integrity of cilia by regulating the abundance of NDE1 and TALPID3. Other yet unidentified candidates also could be playing important roles. Upregulation of NDE1 in *Fbxw7*-null cells (Supplementary Fig. 4) had a dominant effect on ciliary length over TALPID3, leading to an overall reduced number of cells with cilia and a reduction of ciliary length in remaining ciliated cells. Reduced ciliation impacts on Hedgehog activity limiting its activity and resulting in a suppression of differentiation. In *Nde1*-null cells, while ciliation levels are normal, if not enhanced, Hedgehog activity and specifically, GLI2 activity is reduced because of a direct effect of NDE1 on GLI2 protein and activity. Overexpression of GLI2, but not GLI3 in *Nde1*-null cells rescued differentiation. In double mutant cells, the partial rescue in Hedgehog activity could be attributed to an activation of GLI2 triggered by a possible stabilization of TALPID3 levels, due to the loss of FBW7. This increase in TALPID3 could further promote ciliogenesis in the absence of NDE1 and increase Hedgehog activity directly, as has been shown previously by others ^28^ or indirectly. This model is consistent with our data unraveling an interplay between cilia structure and cilia function. Knowledge obtained from our study could have important implications in conditions where Hedgehog activity is upregulated such as in tumors.

FBW7 is one most commonly mutated tumor suppressors. Tumorigenicity induced by the inactivation or loss-of-function mutations in *FBW7* depends on cell context and relevance of target protein(s) ^54^. Based on our current and previous data showing that depletion of FBW7 affects structure and function of primary cilia ^13^, it is tempting to speculate that part of the tumorigenic program induced upon the loss of functional FBW7 could be attributed to cilia and cilia-based signaling. We envision that loss of functional FBW7 could promote tumorigenicity in GLI2-dependent tumors or tumors that arise independently of Hedgehog signaling and they do not require an intact cilium for initiation and/or progression. However, loss or inactivation of *Fbxw7* could have an antitumor or protective effect in Hedgehog-dependent tumors including tumors dependent on inactivating mutations in *PTCH1* or activating mutations in *SMO*, which both require an intact cilium for tumor growth. Inactivating mutations in *PTCH1* and activating mutations in *SMO* have been detected in 73% or 20%, respectively, of patients with basal cell carcinoma (BCC), a stem cell-based malignancy accounting for ∼90% of all solid tumors. Five percent of BCC patients in this study also had inactivating mutations in *FBXW7* ^55^. It would be interesting to know whether tumor aggressiveness correlated with the presence or absence of *PTCH1* or *SMO* with *FBXW7* mutations. According to our model, patients with both mutations (*PTC*H and *FBXW7* or *SMO* and *FBW7*) should have better prognosis.

*NDE1* is mutated in patients with primary microcephaly. We had proposed earlier that loss of NDE1 in radial glial progenitors may have caused a delay in G1 to S transition due to abnormally long cilia ^25^. This hypothesis was supported by *in vivo* experiments in rats, in which NDE1 was depleted *in utero* ^38^. Our current data raise the possibility that reduced Hedgehog activity could also contribute to the delay in cell cycle progression in this context, leading to premature cell cycle exit and depletion or reduction of the radial glial progenitor pool size.

Interestingly, bi-allelic loss-of-function variations in *SMO* or Hedgehog acyl-transferase (*HHAT*) genes in humans led to many developmental anomalies including microcephaly ^56, 57^. While it is unknown whether microcephaly in humans caused by inactivating mutations in *NDE1* is due to abnormally long cilia and/or reduced Hedgehog activity (GLI2), our data lend support to this idea. We attributed the delay in G1-to-S transition in NIH 3T3 and hTERT-RPE1 cells to abnormally long cilia and the extra time needed to disassemble these cilia before progressing into S. The present data lead us to refine this hypothesis to include the possibility that the reduced rate of serum-induced G1/S entry could be attributed to attenuated GLI2 activity in *Nde1*-null cells. Whether this is a specific effect of NDE1 on GLI2 or a general property of all or a subset of negative regulators of ciliogenesis is a subject of future studies.

Joubert syndrome represents a genetically and phenotypically heterogeneous group of disorders characterized by hypoplasia of the cerebellar vermis and other neurologic symptoms. Additional features include skeletal abnormalities, retinal dystrophy, and renal anomalies. TALPID3 is mutated in patients with Joubert syndrome 23, which specifically show hydrocephaly, short-rib thoracic dysplasia 14 with polydactyly, and other skeletal malformations ^46^. Molecular studies in model cell culture systems have shown that TALPID3 is a mother centriolar protein, required for ciliogenesis, and positively regulates Hedgehog signaling activity. Several studies have shown that TALPID3 is required for basal body docking at early stages of ciliogenesis ^26, 27^. However, the specific cell type(s) mediating these *in vivo* effects have not been identified. Our data show that TALPID3 could be a possible target of FBW7 along with NDE1. However, the net effect on cilia in cells lacking *Fbxw7* is a reduction in the number of ciliated cells and reduced ciliary length in remaining ciliated cells, suggesting that NDE1 has a dominant effect on ciliogenesis. From a cilia-centric point of view, it appears that Joubert syndrome 23 and microcephaly may be mediated by diametrically opposite mechanisms, where Joubert Syndrome 23 is caused by loss of cilia, while microcephaly may be caused by abnormally long cilia. However, our data indicate that they are both caused by the failure to couple cilia structure with function, as both diseases seem to share a common defect to maintain a certain level of Hedgehog activity despite cilia being structurally very different.

Finally, our data show that mechanisms mediating ciliary length are relevant in progenitor/stem cell differentiation. Our experiments employing time-dependent deletions of *Fbxw7* before or after commitment to osteoblast differentiation suggest that it may primarily function at the stage which stem cells commit to a certain lineage. This observation was further supported by the positive effect of *Fbxw7* deletion on adipogenesis. Interestingly, RNAi-mediated depletion of FBW7 in the myoblastic stem cell-like line, C2C12 had a positive effect on osteoblast differentiation in response to Bone Morphogenetic Protein 2 (BMP2) treatment ^58^. The different role of FBW7 in osteoblast formation in these cells versus ours can be explained by different routes of osteoblast differentiation. For example, FBW7 may have a negative role in the differentiation of a myoblast-type progenitor to osteoblast, while a positive effect on the differentiation of an MSC to an osteoblast. Despite differences in differentiation mechanisms of different stem-like lines *in vitro*, our mouse data support the positive effect of FBW7 on osteoblast differentiation. We show that FBW7 had an essential role in MSC differentiation to osteoblasts *ex vivo* and postnatal deletion of *Fbxw7* altered bone architecture in 12-week-old male mice. Osteoblast, but not osteoclast, metabolic activity is reduced in these mice. These data lead us to suggest that FBW7 can have an essential role in MSC differentiation to osteoblasts, at least in the animal model used in our studies. Previous studies in MSCs showed that deletion of *Fbxw7* led to an upregulation of NOTCH2, which led to the increased production of the macrophage CCL2 cytokine that promoted metastasis ^59^. No effects on osteoblast differentiation using *ex vivo* cultures or bone architecture were determined in these mice. It is conceivable that FBW7 can have multiple effects on MSCs affecting both differentiation and cytokine production and additional indirect effects on other cell types such as hematopoietic stem cells cannot be excluded.

Overall, our study identified a protein-protein interaction network with an essential role in coupling cilia structure with function. Although the studies were performed in a model system of stem cell differentiation, which is highly amenable to *in vitro, ex vivo*, and *in vivo* approaches of cell differentiation, and in which primary cilia have an established role, new mechanistic information was generated that could help understand cilia function not only in normal conditions, such as stem cell differentiation but also in diverse diseases such as cancer, microcephaly, and Joubert syndrome.

## Methods

### Cell Culture

HEK293T and C3H10T1/2 cells were obtained from ATCC (C3H10T1/2, Clone 8, CCL-226™). HEK293T cells were maintained in Dulbecco’s modified Eagle’s medium (DMEM) supplemented with 10% fetal bovine serum. C3H10T1/2 cells were cultured with Eagle’s Basal Medium (BME) plus 10% heat inactivated fetal bovine serum. Mesenchymal stem cells (MSCs) isolated from the bone marrow of mice, as reported previously ^60^, were cultured with a-MEM medium supplemented with 15% heat inactivated embryonic stem cell qualified fetal bovine serum, 0.22 gr NaHCO3/100 ml media and 1% Penicillin/ Streptomycin/ L-Glutamine (P/S/G, Corning Product Number:30-002-CI). Mouse Embryonic Fibroblasts (MEFS) were maintained in Dulbecco’s Modified Eagle’s Medium (DMEM) plus 10% heat inactivated fetal bovine serum, 1% P/S/G and 1% MEM Nonessential Amino Acids (Corning cellgro, #25-025-CI). MSCs were serum starved with a-MEM medium supplemented with 0.5% heat inactivated embryonic stem cell qualified fetal bovine serum, 0.22gr NaHCO3/100ml media and 1%P/S/G, whereas C3H10T1/2 cells and MEFs were serum starved with reduced serum medium OPTI-MEM I (1X) (Gibco, 31985070). For osteoblast differentiation, C3H10T1/2 cells and MSCs were treated with Mesencult ™ Osteogenic Stimulatory Kit (STEMCELL TECHNOLOGIES, #05504).

Furthermore, MSCs were treated with Mesencult ™ Adipogenic Differentiation Kit (STEMCELL TECHNOLOGIES, #05507) in order to induce adipogenic differentiation. The isolation of MSCs from the bone marrow of 4 weeks old mice was performed based on an established protocol for the isolation and culture of these cells ^60^.

Mouse embryonic fibroblast cultures were prepared as follows: Pregnant females were euthanized when embryos were 13.5-14.5 days old. The uterine horns containing embryos were removed from the mouse and placed in a dish contained sterile PBS on ice. Then the dish was transferred to a new dish that contained PBS in the hood. The uterine wall was teared opened and embryos were moved to 6-well dishes contained sterile PBS. The head of the embryo was removed and used for genotyping. The red tissue was removed in the body cavity and discarded. The remainder of the embryo was used to make MEFS. Each embryo was minced with forceps and transferred to a 15ml conical flasks containing 5ml of trypsin. Furthermore, the cells got into suspension by pipetting up and down and the flask stayed in 37°C incubator for 5 minutes. The cell suspension was transferred in another tube with pre warmed MEF medium and the aforementioned process was repeated 4 times in order to resuspend all the cells. Then the cells were centrifuged and the pellet was re-suspended in 25ml media and plate in a 150mm plate.

### Reagents

Ciliobrevin A was purchased from Tocris (#4529) and MLN4924 (Pevonedistat, # HY-70062) was purchased from MedChemExpress (MCE). Paraformaldehyde (#J19943-K2) and Prolonged Diamond DAPI (#P36966) were purchased from Thermofisher. Finally, 4-hydroxytamoxifen was purchased from Sigma (#H6278) and SAG was purchased from Abcam (#ab142160). The working concentration of: a) ciliobrevin A is 50µM, b) SAG is 1µg/ml, c) 4-hydroxytamoxifen for the ex vivo experiments is 2µg/ml and D) MLN4924 is 0.25 or 0.5μM.

### Plasmids

hNde1 cDNA was obtained from Open Biosystems and was subcloned into a pFLAG-CMV-2 vector (Kim et al. 2011). Myc-GLI2, HA-GLI3P1-4A-FLAG, HA-GLI3P1-6A-FLAG, Myc-GLI2 delta N, MYC-FBW7 plasmid were obtained from Addgene. TALPID3-DDK-MYC was obtained from ORIGENE. FBW7α-HA, FBW7β-HA, FBW7γ-HA, FBW7α-HA^ΔF^, FBW7α-HA^R456C^ were designed by having a 4xHA tag at the C-terminus. Gli-BS plasmid was a gift from Dr. Brad Yoder (UAB, AL).

### CRISPR–CAS9 knockout

C3H10T1/2^*Fbxw7KO*^ and C3H10T1/2^*Nde1KO*^ clones were generated as follows: C3H10T1/2 cells were transfected with a lentiviral vector encoding Cas9, resistance to puromycin and single guide RNA (sgRNA) sequences specific for *Fbxw7* (5’-CACCGATGAAGTCTCGCTGGAACTG-3’) or *Nde1* (5’-CACCGACTCCAGCTCCATGCGAAGG -3’). After transfection, the cells underwent serial dilutions and were plated under puromycin selection (1 ug/ml) for 3 weeks. After selection, single colonies were extracted and grown. For the generation of the C3H10T1/2^*Fbxw7KO*;*Nde1KO*^ clone, a C3H10T1/2^*Fbxw7*KO^ clone was co-transfected with the aforementioned lentiviral vector encoding Cas9, resistance to puromycin and a single guide RNA (sgRNA) sequence specific for *Nde1* (5’-CACCGACTCCAGCTCCATGCGAAGG -3’) and with an empty vector encoding resistance to G418 in order to be able to select C3H10T1/2^*Fbxw7KO*;*Nde1KO*^clones. The G418 concertation that was used for the selection of the double KO clones was 1 mg/ml. For the generation of C3H10T1/^*Ift88KO*^ clones the same methodology was applied: C3H10T1/2 cells were transfected with a lentiviral vector encoding Cas9, resistance to puromycin and single guide RNA (sgRNA) sequences specific for *IFT88* (5’-CACCGCAACCCAGCCTATGATACTG-3’) The resulting clones were evaluated for deletion of the genes of interest by DNA sequencing or/and Western blotting.

### Transient transfection

siRNAs were transfected into C3H10T1/2 cells using RNAiMAX reagent (Invitrogen) or using Lipofectamine 2000 (Invitrogen) if co-transfected with plasmids according to the manufacturer’s instructions. Transfections with plasmids in C3H10T1/2 were done by using Lipofectamine LTX with PLUS™ Reagent (Invitrogen). Transient transfections were done in 293T cells by using calcium phosphate method, as previously described (Kim *et al*, 2011).

### siRNA sequences

Mouse *Fbxw7* -specific smart pool siRNA was obtained from Dharmacon (#L-041553-01-0005). Non-targeting siRNA pool was obtained from Dharmacon (#D-001810-10-05). Mouse *Talpid3* - specific smart pool siRNA was obtained from Dharmacon (#L-043740-01-0005).

### Immunoblotting

HEK293T cells, wild type C3H10T1/2 cells, C3H10T1/2^*Fbxw7*KO^, C3H10T1/2^*Nde1KO*^ and C3H10T1/2 ^*Fbxw7KO;Nde1KO*^ clones were lysed in 1% Triton X-100, 150mM NaCl, 10 mM Tris-HCl at pH 7.5, 1mM EGTA, 1mM EDTA, 10% sucrose and a protease inhibitor cocktail (Roche Applied Science), phosphatase inhibitor cocktail (PhosSTOP EASYpack, ROCHE) at 4C for 30 min. Cell lysates were separated with SDS-PAGE. Antibodies were used against ALP (Thermofisher, 1:200), NDE1 (Proteintech, 1:1,000), β-tubulin (Santa Cruz, 1:1,000), TALPID3 (Proteintech 1:1000), HA (Santacruz 1:1000), MYC-Tag (Cell Signaling), FLAG (Sigma 1:1000), FBW7 (Bethyl Laboratories. A301-720A; A301-721A 1:500), NDEL1 (Proteintech, 1:1000), GAPDH (GeneTex, 1:2000) and IFT88 (Proteintech, 1:000). Densitometric quantification was performed with the Licor Image Studio software.

### Immunoprecipitation

Wild type C3H10T1/2, *Fbxw7*-null C3H10T1/2, *Nde1-null* C3H10T1/2 or double mutant cell lysates were incubated with α-FBW7 antibody (Abnova) overnight to immunoprecipitate FBW7. The antigen-antibody complexes were then incubated with Protein G Sepharose beads for 3 hours in 4°C and were analyzed for the presence of FBW7 by Western blot. The same immunoprecipitation conditions were applied for the interactions between GLI2-NDE1, or TALPID3 and FBW7. In the GLI2-NDE1 interactions cell lysates were incubated with the FLAG antibody, and in the TALPID3-FBW7 interaction cell lysates were incubated either with a-TALPID3 or with a-HA.

### Indirect immunofluorescence

MSCs, Mouse embryonic fibroblasts and C3H10T1/2 cells were grown on glass coverslips and fixed in 4% paraformaldehyde, permeabilized in 0.1% Triton X-100 in PBS, blocked in 3% heat-activated goat serum (or donkey serum for ALP) /0.1% Triton X-100 in PBS (blocking buffer), and incubated overnight with primary antibodies diluted in blocking buffer at 4°C. Primary antibodies were used against mouse acetylated a-tubulin at 1:1,000 (Sigma Aldrich, 1:1000), CD106 (Abcam, 1:200) or goat ALP (Thermofisher, 1:50). Cells were washed three times with PBS and incubated for 2 h at 4°C with appropriate combinations of AlexaFluor-conjugated secondary antibodies (Invitrogen,1:2000) for 2 hours at 4C protected from light. Excess of secondary antibodies were removed by four washes in PBS Samples were mounted with Diamond DAPI (Thermofisher) to counterstain the nuclei. Images were obtained with an Olympus FV1000 confocal microscope and processed with ImageJ software for ciliary length measurements.

### Quantitative polymerase chain reaction (qPCR)

RNA was extracted and purified from Wild type mesenchymal stem cells or *Fbxw7* null-MSCs at different time points of osteogenic treatment or at 5days of adipogenic treatment using Trizol reagent (Invitrogen). RNA was reverse-transcribed to cDNA and samples were amplified by qPCR. mRNA levels of the genes of interest were normalized to wild type via the ΔΔCt method. Primers used for qPCR were *Fbxw7* Fw: 5’-ACTGGAGAATTTTGGCTGAGGAT-3, *Fbxw7* Rv: 5’-ATGGGCTGTGTATGAAACCTGG-3’, *Runx2* Fw: 5’-*CCGAAATGCCTCCGCTGTTA*-3’, *Runx2* Rv: 5’-*TGAAACTCTTGCCTCGTCCG*-3’, *OSX* Fw: 5’-*GATGGCGTCCTCTCTGCTTGA*-3’, *OSX* Rv: 5’-*CAGGGTTGTTGAGTCCCGCA*-3’, *ALP* Fw: 5’-*GCAAGGACATCGCATATCAGC*-3’, ALP Rv: 5’-*TCCAGTTCGTATTCCACATCAGT*-3’, *OCN* Fw: 5’-*AGCGGCCCTGAGTCTG*-3’, *OCN* Rv: 5’-*CTGGGCTGGGGACTGA*-3’, *Optn Fw: AGCTTGGCTTATGGACTGAGG, Optn* Rv: *AGACTCACCGCTCTTCATGTG, GAPDH:* Fw: 5’-*AAAATGGTGAAGGTCGGTGTG*-3’, *GAPDH:* Rv: 5’-*AATGAAGGGGTCGTTGATGG*-3’, *CEBP1a:* Fw: 5’-*GGGAACGCAACAACATCGC*-3’, *CEBP1a:* Rv: 5’-*GCGGTCATTGTCACTGGTCA*-3’, *Adiponectin:* Fw: 5’-*GCAGAGATGGCACTCCTGGA*-3’, *Adiponectin:* Rv: 5’-*CCCTTCAGCTCCTGTCATTCC*-3’, *PPAR*γ: Fw: 5’-*GTGGGGATAAAGCATCAGGC*-3’, *PPAR*γ: Rv: 5’-*TCCGGCAGTTAAGATCACACC*-3’, *TBP:* Fw: 5’-*TCTACCGTGAATCTTGGCTGT*-3’, *TBP:* Rv: 5’-*GTCCGTGGCTCTCTTATTCTCA*-3’.

### Osteogenic differentiation

Osteogenic differentiation induction was performed as follows: Mesenchymal stem cells or C3H10T1/2 cells were trypsinized and seeded in a 96-well or 24-well plate respectively. When 100 % confluency was reached, the medium was replaced with osteogenic induction medium (STEMCELL TECHNOLOGIES, #05504). The osteogenic differentiation for mesenchymal stem cells was evaluated by Alizarin Red S staining at days 14 and 21 and 28 days of differentiation and quantitative real-time PCR of various osteogenic marker genes at these days. Alkaline phosphatase (ALP) staining was used as an indicator of osteoblast differentiation in C3H10T1/2 at 5 days of osteogenic treatment. Triplicate tests were conducted in each experiment.

### Alizarin Red S Staining

Mesenchymal stem cells were differentiated to osteoblasts with the osteogenic induction medium. Then, cells were washed with PBS, fixed with 4% paraformaldehyde and incubate at room temperature for 15minutes. The fixative was then removed and cells were washed twice with PBS. Then samples were incubated with Alizarin Red Solution at room temperature for 20minutes. After 20 minutes of incubation, the dye was removed and the cells were gently washed three times with ddH20. After the last wash plates were let to dry and were ready for image acquisition at the dissection microscope.

### Alkaline Phosphatase Staining

Undifferentiated C3H10T1/2 cells show weak alkaline phosphatase (ALP) activity whereas differentiated osteoblasts show very high activity of ALP. ALP activity is therefore a good indicator of osteoblast differentiation. One BCIP/NBT tablet (SigmaFast™ BCIP-NBT, Sigma Aldrich) was dissolved in 10ml distilled ddH20 to prepare the substrate solution which detects ALP by staining the cells blue-violet when ALP is present. It should be stored in the dark and used within 2hours. The washing buffer was made by adding 0.05% Tween 20 to PBS without Ca^++/^Mg^++^ (Corning, #21-040-CM). Cells were washed one time with PBS and fixed with 4% paraformaldehyde for 60 seconds. After the fixation, cells were washed with washing buffer and then treated with BCIP/NBT substrate solution at room temperature in the dark for 10 minutes. The staining process is being observed every 2-3 minutes. Cells are then washed with 1x with washing buffer and 1x with PBS respectively. After the last wash, the staining results were analyzed at the dissection microscope.

### Adipogenic differentiation

Adipogenic differentiation induction was performed as follows: Mesenchymal stem cells were trypsinized and seeded in a 96-well plate respectively. When 100 % confluency was reached, the medium was replaced with Mesencult ™ Adipogenic Differentiation Kit (STEMCELL TECHNOLOGIES, #05507). The adipogenic differentiation for mesenchymal stem cells was evaluated by Oil Red O staining and quantitative real-time PCR of adipogenic marker genes at 5 days of differentiation.

### Oil O Red Staining

Oil Red O was purchased from Sigma (#O-0625). The stock solution was made by adding 0.35g Oil Red O in 100ml of isopropanol. The Oil Red O working solution was made by mixing 6ml of Oil Red O stock solution with 4ml of ddH20, let it at room temperature for 20minutes followed by filtering. Cells were washed 1X with PBS and fixed with 4% of paraformaldehyde for 15 minutes at room temperature. Cells were washed twice with ddH20 and then 1X with 60% isopropanol for 5 minutes at RT. After the aspiration of isopropanol cells should be dried completely and then incubated with Oil Red O working solution for 10minutes at room temperature. The next step is the washing of cells 4X with ddH20 and the acquisition of the images acquisition of the images at the dissection microscope.

### Hedgehog Activity

Hedgehog Activity was analyzed in C3H10T1/2 or HEK293T cells by using a GLIBS reporter plasmid, expressing firefly luciferase in a manner dependent on the activity of Hedgehog pathway. Full growth medium was replaced 24h after transfection and cells were lysed in lysis buffer (Promega) 48h after transfection. Cell lysates were incubated with Stop & Glo substrates (Promega) and luciferase activity was measured in a Synergy Neo2 reader (Biotek).

### Ubiquitinylation assay

HEK293T cells were transiently co-transfected with TALPID3-DDK-MYC and with or pcB6-His ubiquitin (provided by R. Baer), FBW7α-HA, FBW7α-HA^ΔF^, FBW7α-HA^R456C^. *In vivo* ubiquitinylation assay was done according to Dipak et al (2015).

### Micro-Computed Tomography (µCT) Analysis

All procedures were conducted based on the requirements of the IACUC. In this study, *UbcCre*^*ERT2*^; *Fbxw7*^*f/f*^ male mice were crossed with *Fbxw7*^*f/f*^ female mice (all mice have C57Bl/6-background). For the deletion of *Fbxw7*, nursing dams were intraperitoneally injected with 4-Hydroxy-Tamoxifen from postnatal days 2 to 6 (P2-P6) as previously described ^61^. Then mice at 12 weeks old were sacrificed, soft tissue was removed and femur was analyzed at the UCSF CCMBM Skeletal Biology and Biomechanics Core for changes in 3-dimension structural parameters.

### ELISA

Serum was obtained from 12-week-old mice and analyzed by ELISA for Mouse Bone Specific alkaline phosphatase (BALP, MyBiosource, #MBS281206) and Mouse Tartrate-Resistant Acid Phosphatase 5b (TRACP-5b, MyBiosource, #MBS701767) levels according to the manufacturer’s protocols.

### Statistics

The statistical analyses were performed by Software GraphPad Prism 8.3. Student’s t test or one-way ANOVA were performed for the quantification analysis of the results based on the number of comparison groups, followed by the proper *ad hoc* test.

## Author contributions

EP and VG performed experiments. NS performed microcomputing analysis. EP, WC, MBH, and LT analyzed data. EP and LT wrote the manuscript.

## Acknowledgments

We thank Dr. Lorin Olson for comments. This work was supported by AR064211 (LT and MBH), DK59599 (LT), and DK12165601 (LT and WC) from NIH; VA-BLR&D I01BX003453 (WC) from VA, Presbyterian Health Foundation (MBH), and Oklahoma Center for Adult Stem Cell Research (LT).

## References

1. Malicki, J.J. & Johnson, C.A. The Cilium: Cellular Antenna and Central Processing Unit. Trends Cell Biol 27, 126–140 (2017).

2. Anvarian, Z., Mykytyn, K., Mukhopadhyay, S., Pedersen, L.B. & Christensen, S.T. Cellular signalling by primary cilia in development, organ function and disease. Nat Rev Nephrol 15, 199–219 (2019).

3. Berbari, N.F., O’Connor, A.K., Haycraft, C.J. & Yoder, B.K. The primary cilium as a complex signaling center. Curr Biol 19, R526–535 (2009).

4. Hilgendorf, K.I., Johnson, C.T. & Jackson, P.K. The primary cilium as a cellular receiver: organizing ciliary GPCR signaling. Curr Opin Cell Biol 39, 84–92 (2016).

5. Bangs, F. & Anderson, K.V. Primary Cilia and Mammalian Hedgehog Signaling. Cold Spring Harb Perspect Biol 9 (2017).

6. Mukhopadhyay, S. & Rohatgi, R. G-protein-coupled receptors, Hedgehog signaling and primary cilia. Semin Cell Dev Biol 33, 63–72 (2014).

7. Liu, H., Kiseleva, A.A. & Golemis, E.A. Ciliary signalling in cancer. Nature reviews. Cancer 18, 511–524 (2018).

8. Mirvis, M., Siemers, K.A., Nelson, W.J. & Stearns, T.P. Primary cilium loss in mammalian cells occurs predominantly by whole-cilium shedding. PLoS biology 17, e3000381 (2019).

9. Mirvis, M., Stearns, T. & James Nelson, W. Cilium structure, assembly, and disassembly regulated by the cytoskeleton. Biochem J 475, 2329–2353 (2018).

10. Pugacheva, E.N., Jablonski, S.A., Hartman, T.R., Henske, E.P. & Golemis, E.A. HEF1-dependent Aurora A activation induces disassembly of the primary cilium. Cell 129, 1351–1363 (2007).

11. Sanchez, I. & Dynlacht, B.D. Cilium assembly and disassembly. Nat Cell Biol 18, 711–717 (2016).

12. Wang, L. & Dynlacht, B.D. The regulation of cilium assembly and disassembly in development and disease. Development 145 (2018).

13. Maskey, D. et al. Cell cycle-dependent ubiquitylation and destruction of NDE1 by CDK5-FBW7 regulates ciliary length. Embo j 34, 2424–2440 (2015).

14. Nikonova, A.S. & Golemis, E.A. The tumor suppressor FBW7 controls ciliary length. Embo j 34, 2388–2390 (2015).

15. Huangfu, D. et al. Hedgehog signalling in the mouse requires intraflagellar transport proteins. Nature 426, 83–87. (2003).

16. Haycraft, C.J. et al. Gli2 and Gli3 localize to cilia and require the intraflagellar transport protein polaris for processing and function. PLoS Genet 1, e53. Epub 2005 Oct 2028. (2005).

17. Niewiadomski, P. et al. Gli protein activity is controlled by multisite phosphorylation in vertebrate Hedgehog signaling. Cell reports 6, 168–181 (2014).

18. Briscoe, J. & Therond, P.P. The mechanisms of Hedgehog signalling and its roles in development and disease. Nat Rev Mol Cell Biol 14, 416–429 (2013).

19. Caplan, A.I. Mesenchymal stem cells. J Orthop Res 9, 641–650 (1991).

20. Mendez-Ferrer, S. et al. Mesenchymal and haematopoietic stem cells form a unique bone marrow niche. Nature 466, 829–834 (2010).

21. Hass, R., Kasper, C., Bohm, S. & Jacobs, R. Different populations and sources of human mesenchymal stem cells (MSC): A comparison of adult and neonatal tissue-derived MSC. Cell Commun Signal 9, 12 (2011).

22. Hoey, D.A., Tormey, S., Ramcharan, S., O’Brien, F.J. & Jacobs, C.R. Primary Cilia-Mediated Mechanotransduction in Human Mesenchymal Stem Cells. STEM CELLS 30, 2561–2570 (2012).

23. Takeishi, S. & Nakayama, K.I. Role of Fbxw7 in the maintenance of normal stem cells and cancer-initiating cells. British Journal of Cancer 111, 1054–1059 (2014).

24. Welcker, M. & Clurman, B.E. FBW7 ubiquitin ligase: a tumour suppressor at the crossroads of cell division, growth and differentiation. Nature reviews. Cancer 8, 83–93 (2008).

25. Kim, S. et al. Nde1-mediated inhibition of ciliogenesis affects cell cycle re-entry. Nature Cell Biology 13, 351–360 (2011).

26. Davey, M.G. et al. The chicken talpid3 gene encodes a novel protein essential for Hedgehog signaling. Genes Dev 20, 1365–1377 (2006).

27. Kobayashi, T., Kim, S., Lin, Y.C., Inoue, T. & Dynlacht, B.D. The CP110-interacting proteins Talpid3 and Cep290 play overlapping and distinct roles in cilia assembly. J Cell Biol 204, 215–229 (2014).

28. Li, J. et al. PKA-mediated Gli2 and Gli3 phosphorylation is inhibited by Hedgehog signaling in cilia and reduced in Talpid3 mutant. Dev Biol 429, 147–157 (2017).

29. Yin, Y. et al. The Talpid3 gene (KIAA0586) encodes a centrosomal protein that is essential for primary cilia formation. Development 136, 655–664 (2009).

30. Tetzlaff, M.T. et al. Defective cardiovascular development and elevated cyclin E and Notch proteins in mice lacking the Fbw7 F-box protein. Proc Natl Acad Sci U S A 101, 3338–3345 (2004).

31. Baddoo, M. et al. Characterization of mesenchymal stem cells isolated from murine bone marrow by negative selection. J Cell Biochem 89, 1235–1249 (2003).

32. Yang, Z.X. et al. CD106 identifies a subpopulation of mesenchymal stem cells with unique immunomodulatory properties. PloS one 8, e59354 (2013).

33. Chou, D.B. et al. Stromal-derived IL-6 alters the balance of myeloerythroid progenitors during Toxoplasma gondii infection. Journal of leukocyte biology 92, 123–131 (2012).

34. Peister, A. et al. Adult stem cells from bone marrow (MSCs) isolated from different strains of inbred mice vary in surface epitopes, rates of proliferation, and differentiation potential. Blood 103, 1662–1668 (2004).

35. Zhao, L., Li, G., Chan, K.M., Wang, Y. & Tang, P.F. Comparison of multipotent differentiation potentials of murine primary bone marrow stromal cells and mesenchymal stem cell line C3H10T1/2. Calcified tissue international 84, 56–64 (2009).

36. Katagiri, T. et al. The non-osteogenic mouse pluripotent cell line, C3H10T1/2, is induced to differentiate into osteoblastic cells by recombinant human bone morphogenetic protein-2. Biochem Biophys Res Commun 172, 295–299 (1990).

37. Inaba, H. et al. Ndel1 suppresses ciliogenesis in proliferating cells by regulating the trichoplein-Aurora A pathway. J Cell Biol 212, 409–423 (2016).

38. Doobin, D.J., Kemal, S., Dantas, T.J. & Vallee, R.B. Severe NDE1-mediated microcephaly results from neural progenitor cell cycle arrests at multiple specific stages. Nature communications 7, 12551 (2016).

39. Sasaki, S. et al. Complete loss of Ndel1 results in neuronal migration defects and early embryonic lethality. Mol Cell Biol 25, 7812–7827 (2005).

40. Nakamura, T. et al. Novel hedgehog agonists promote osteoblast differentiation in mesenchymal stem cells. J Cell Physiol 230, 922–929 (2015).

41. Nakamura, T. et al. Induction of osteogenic differentiation by hedgehog proteins. Biochem Biophys Res Commun 237, 465–469 (1997).

42. Firestone, A.J. et al. Small-molecule inhibitors of the AAA+ ATPase motor cytoplasmic dynein. Nature 484, 125–129 (2012).

43. Zhang, Q. et al. FBXW7 Facilitates Nonhomologous End-Joining via K63-Linked Polyubiquitylation of XRCC4. Mol Cell 61, 419–433 (2016).

44. Roessler, E. et al. A previously unidentified amino-terminal domain regulates transcriptional activity of wild-type and disease-associated human GLI2. Hum Mol Genet 14, 2181–2188 (2005).

45. Schulein-Volk, C. et al. Dual regulation of Fbw7 function and oncogenic transformation by Usp28. Cell reports 9, 1099–1109 (2014).

46. Stephen, L.A. et al. TALPID3 controls centrosome and cell polarity and the human ortholog KIAA0586 is mutated in Joubert syndrome (JBTS23). eLife 4 (2015).

47. Goto, H., Inaba, H. & Inagaki, M. Mechanisms of ciliogenesis suppression in dividing cells. Cellular and molecular life sciences : CMLS 74, 881–890 (2017).

48. Higgins, M., Obaidi, I. & McMorrow, T. Primary cilia and their role in cancer. Oncol Lett 17, 3041–3047 (2019).

49. Yang, S. & Wang, C. The intraflagellar transport protein IFT80 is required for cilia formation and osteogenesis. Bone 51, 407–417 (2012).

50. Qiu, N., Cao, L., David, V., Quarles, L.D. & Xiao, Z. Kif3a deficiency reverses the skeletal abnormalities in Pkd1 deficient mice by restoring the balance between osteogenesis and adipogenesis. PloS one 5, e15240 (2010).

51. Corrigan, M.A., Ferradaes, T.M., Riffault, M. & Hoey, D.A. Ciliotherapy Treatments to Enhance Biochemically-and Biophysically-Induced Mesenchymal Stem Cell Osteogenesis: A Comparison Study. Cellular and molecular bioengineering 12, 53–67 (2019).

52. Johnson, G.P. et al. Mesenchymal stem cell mechanotransduction is cAMP dependent and regulated by adenylyl cyclase 6 and the primary cilium. J Cell Sci 131 (2018).

53. Lehti, M.S. et al. Cilia-related protein SPEF2 regulates osteoblast differentiation. Scientific reports 8, 859 (2018).

54. Yumimoto, K. & Nakayama, K.I. Recent insight into the role of FBXW7 as a tumor suppressor. Semin Cancer Biol (2020).

55. Bonilla, X. et al. Genomic analysis identifies new drivers and progression pathways in skin basal cell carcinoma. Nat Genet 48, 398–406 (2016).

56. Le, T.L. et al. Bi-allelic Variations of SMO in Humans Cause a Broad Spectrum of Developmental Anomalies Due to Abnormal Hedgehog Signaling. Am J Hum Genet 106, 779–792 (2020).

57. Abdel-Salam, G.M.H. et al. Biallelic novel missense HHAT variant causes syndromic microcephaly and cerebellar-vermis hypoplasia. Am J Med Genet A 179, 1053–1057 (2019).

58. Yumimoto, K., Matsumoto, M., Onoyama, I., Imaizumi, K. & Nakayama, K.I. F-box and WD repeat domain-containing-7 (Fbxw7) protein targets endoplasmic reticulum-anchored osteogenic and chondrogenic transcriptional factors for degradation. J Biol Chem 288, 28488–28502 (2013).

59. Yumimoto, K. et al. F-box protein FBXW7 inhibits cancer metastasis in a non-cell-autonomous manner. J Clin Invest 125, 621–635 (2015).

60. Huang, S. et al. An improved protocol for isolation and culture of mesenchymal stem cells from mouse bone marrow. J Orthop Translat 3, 26–33 (2015).

61. Gerakopoulos, V., Ngo, P. & Tsiokas, L. Loss of polycystins suppresses deciliation via the activation of the centrosomal integrity pathway. Life Sci Alliance 3 (2020).

