## Supplemental Data for "FBW7 couples structural integrity with functional output of primary cilia"

### Supplementary Figure Legends

**Supplementary Fig. 1. *Fbxw7* deletion after osteogenic induction mildly suppresses differentiation of *UbcCre$^{ERT2}$; Fbxw7$^{f/f}$* MSCs to osteoblasts.**

**a)** Diagram showing experimental setup for ex vivo deletion of *Fbxw7*. Isolation of MSCs from the bone marrow of *UbcCre$^{ERT2}$; Fbxw7$^{f/f}$* mice (1) was followed by 24h osteogenic treatment (2), 24h 4OHT-induced deletion of *Fbxw7* (3), continuation of osteogenic treatment (4) and analysis of osteoblast differentiation (5).

**b-d)** Osteoblast differentiation of *UbcCre$^{ERT2}$; Fbxw7$^{f/f}$*-derived MSCs treated with mock or 4OHT after initial osteogenic induction (n=3 different mice). Differentiation was measured at 14 and 21 days after osteogenic induction via Alizarin Red S staining (b) and mRNA levels of osteoblast differentiation markers *Runx2*, *Osterix* (*Osx*), *Osteocalcin* (Ocn), *Osteopontin* (*Opnt*) and Alkaline phosphatase (*Alp*) at 14 (c) and 21 days (d). mRNA levels of *Fbxw7* were analyzed to confirm deletion of the gene. Data are presented as means ± SEM. Student's t-test, $^{**}p < 0.01$, $^{***}p < 0.001$, $^{****}p < 0.0001$.

**Supplementary Fig. 2. No effect of 4OHT in osteoblast differentiation of wild type MSCs.**

**a)** Osteoblast differentiation of *Fbxw7$^{f/f}$*-derived MSCs treated with mock or 4OHT. Differentiation was measured at 14, 21 and 28 days via Alizarin Red S staining.

**b)** Confirmation of deletion of *Fbxw7* in MSCs of *UbcCre$^{ERT2}$; Fbxw7$^{f/f}$* after various duration of treatment with 4OHT. Black arrow indicates the PCR product after disruption of *Fbxw7*.

**Supplementary Fig. 3. Deletion of *Fbxw7* before adipogenic induction increases adipogenesis in *UbcCre$^{ERT2}$; Fbxw7$^{f/f}$* MSCs.**

**a-j)** Adipogenic differentiation of *UbcCre$^{ERT2}$; Fbxw7$^{f/f}$*-derived MSCs treated with mock or 4OHT before (a-d) (n=3 different mice) or after initial adipogenic induction (e-h) (n=4 different mice). Differentiation was measured via Oil red O staining (a,e) and mRNA levels of adipogenic differentiation markers: *CEBPa* (b,f) *Adiponectin* (c,g) and *Pparγ* (d,h). Data are presented as means ± SEM. Student's t-test, *$p < 0.05$, **$p < 0.01$, ****$p < 0.0001$.

**Supplementary Fig. 4. Validation of deletion or downregulation of indicated target genes in C3H10T1/2.**

**a)** Immunoprecipitation of endogenous FBW7 in wild type, *Fbxw7KO*, *Nde1KO* and *Fbxw7Nde1KO* C3H10T1/2 clones generated via CRISPR-Cas9 gene editing with *Fbxw7-* or *Nde1-* specific sgRNAs. Bottom: Sanger sequencing of exon 1 of mouse *Fbxw7* in *Fbxw7KO* C3H10T1/2 cells revealed a deletion around the Cas9 cleavage site.

**b)** Expression levels of endogenous NDE1 in the *Fbxw7KO* C3H10T1/2 clone under 24h serum starvation conditions. Arrow indicates the band denoting NDE1.

**c)** Expression levels of endogenous NDE1 in various candidate *Nde1KO* clones. *Nde1$^{+/+}$* and *Nde1$^{-/-}$* MEF controls are shown in lanes 1 and 2, respectively, indicating that the lower band of the doublets is NDE1. Bottom: Sanger sequencing of the clone indicated with the red asterisk revealed a deletion around the Cas9 cleavage site.

**d)** Expression levels of endogenous NDE1 in various candidate *Fbxw7Nde1KO* clones. Previously generated *Fbxw7KO* C3H10T1/2 clone was transfected with *Nde1-* specific sgRNA to generate double *Fbxw7Nde1KO* C3H10T1/2 clones. *Nde1*$^{+/+}$ and *Nde1*$^{-/-}$ MEF controls are shown in lanes 13 and 14, respectively, indicating that the lower band in the doublets is NDE1.

**e)** Absence of NDEL1 expression in C3H10T1/2 clones. NDEL1 overexpression in HEK293T cells was used as a positive control.

**f,g)** Expression levels of TALPID3 in wild type (f) or *Fbxw7Nde1KO* (g) C3H10T1/2 cells transfected with a *Talpid3* construct and mock or *Talpid3* specific siRNA

**Supplementary Fig. 5. Disruption of primary cilia compromises osteoblast differentiation in C3H10T1/2 cells.**

**a,b)** ALP staining (a) and expression levels of ALP (b) in C3H10T1/2 cells after treatment with DMSO or Ciliobrevin A and osteogenic induction. 2X indicates magnification of corresponding 1X images.

**c)** Generation of Ift88KO C3H10T1/2 via CRISPR-Cas9 gene editing. Expression levels of endogenous IFT88. Red asterisk indicates null clone selected.

**d)** Wild type or the Ift88KO clone from (c) was induced via osteogenic medium and differentiation was analyzed via ALP staining. 2X indicates magnification of corresponding 1X images.

**e)** Representative ALP staining of wild type and *Nde1KO* C3H10T1/2 cells transfected with the indicated constructs.

**f)** Representative ALP staining of wild type and *Fbxw7KO* C3H10T1/2 cells transfected with the indicated constructs.

**Supplementary Fig. 6. Inhibition of FBW7 by MLN4924 before osteogenic induction severely compromises osteoblast differentiation in C3H10T1/2 cells.**

**a)** ALP staining of wild type C3H10T1/2 cells after the indicated treatments. 2X indicates magnification of corresponding 1X images.

**b,c)** Expression levels of ALP in wild type C3H10T1/2 cells after the indicated treatments (b) and summary data (c). Data are presented as means ± SEM. One-way ANOVA with Dunnett's multiple comparisons test, ****$p < 0.0001$.

Suppl. Fig 1

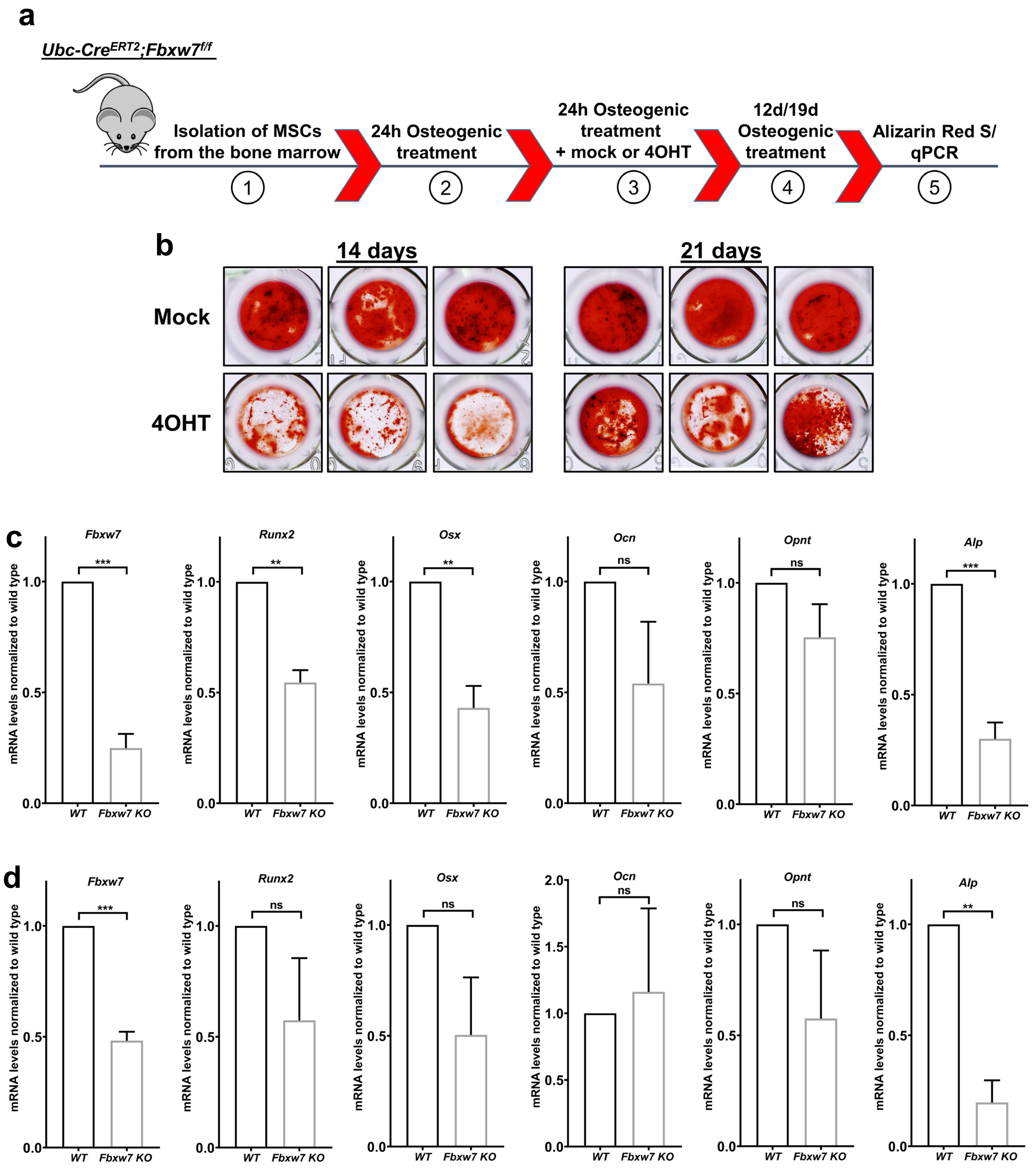

a
Ubc-CreERT2;Fbxw7f/f
Isolation of MSCs from the bone marrow
24h Osteogenic treatment
24h Osteogenic treatment + mock or 4OHT
12d/19d Osteogenic treatment
Alizarin Red S/ qPCR
1
2
3
4
5
b
14 days
21 days
Mock
4OHT
c
Fbxw7
Runx2
Osx
Ocn
Opnt
Alp
***
**
**
ns
ns
***
mRNA levels normalized to wild type
1.0
0.5
0.0
WT
Fbxw7 KO
d
Fbxw7
Runx2
Osx
Ocn
Opnt
Alp
***
ns
ns
ns
ns
**
2.0
1.5
1.0
0.5
0.0
mRNA levels normalized to wild type
WT
Fbxw7 KO

**Suppl. Fig 2**

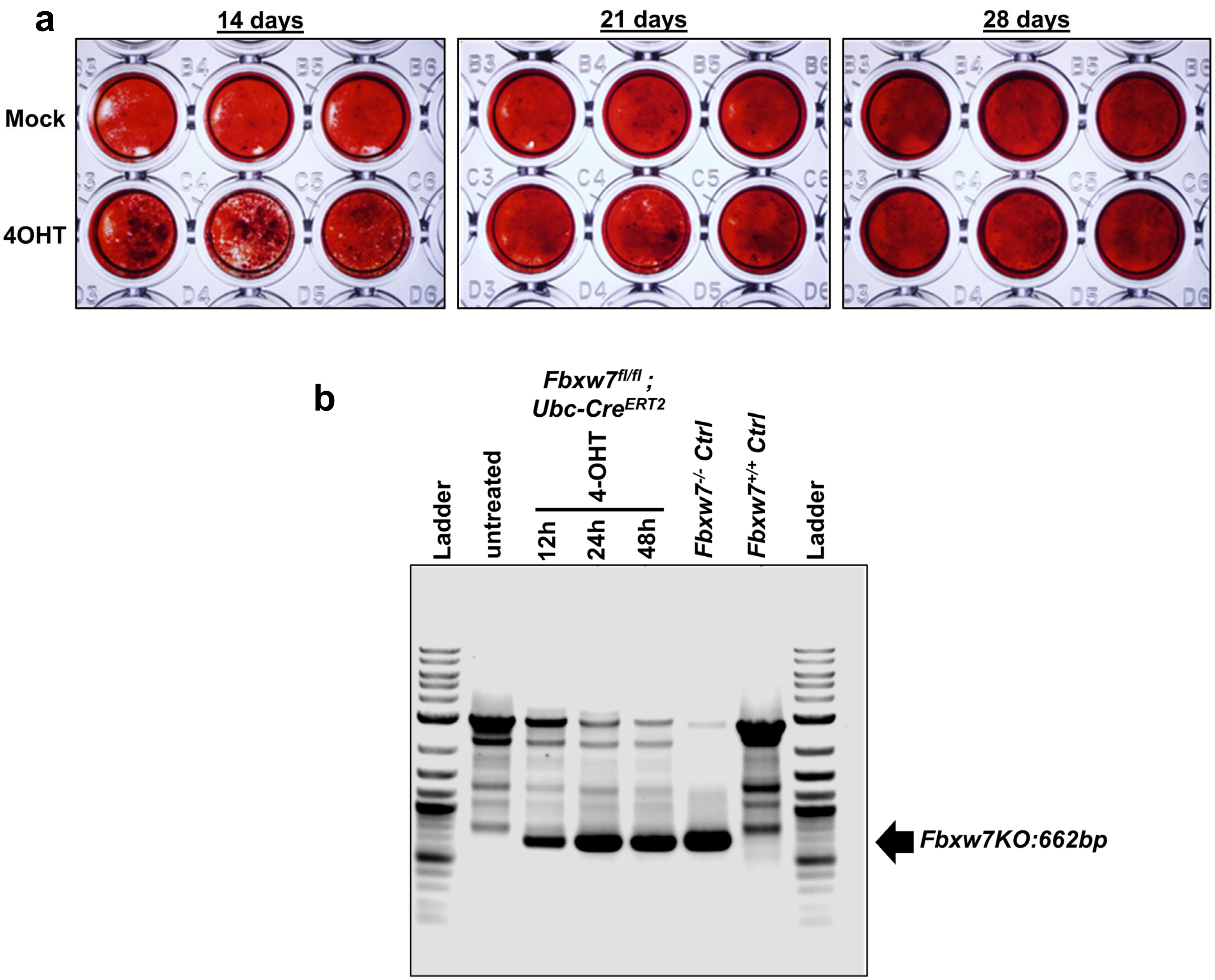
a
14 days
21 days
28 days
Mock
4OHT
B3
B4
B5
B6
C3
C4
C5
C6
D3
D4
D5
D6
b
Fbxw7fl/fl ;
Ubc-CreERT2
Ladder
untreated
4-OHT
12h
24h
48h
Fbxw7-/- Ctrl
Fbxw7+/+ Ctrl
Ladder
Fbxw7KO:662bp

### Suppl. Fig 3

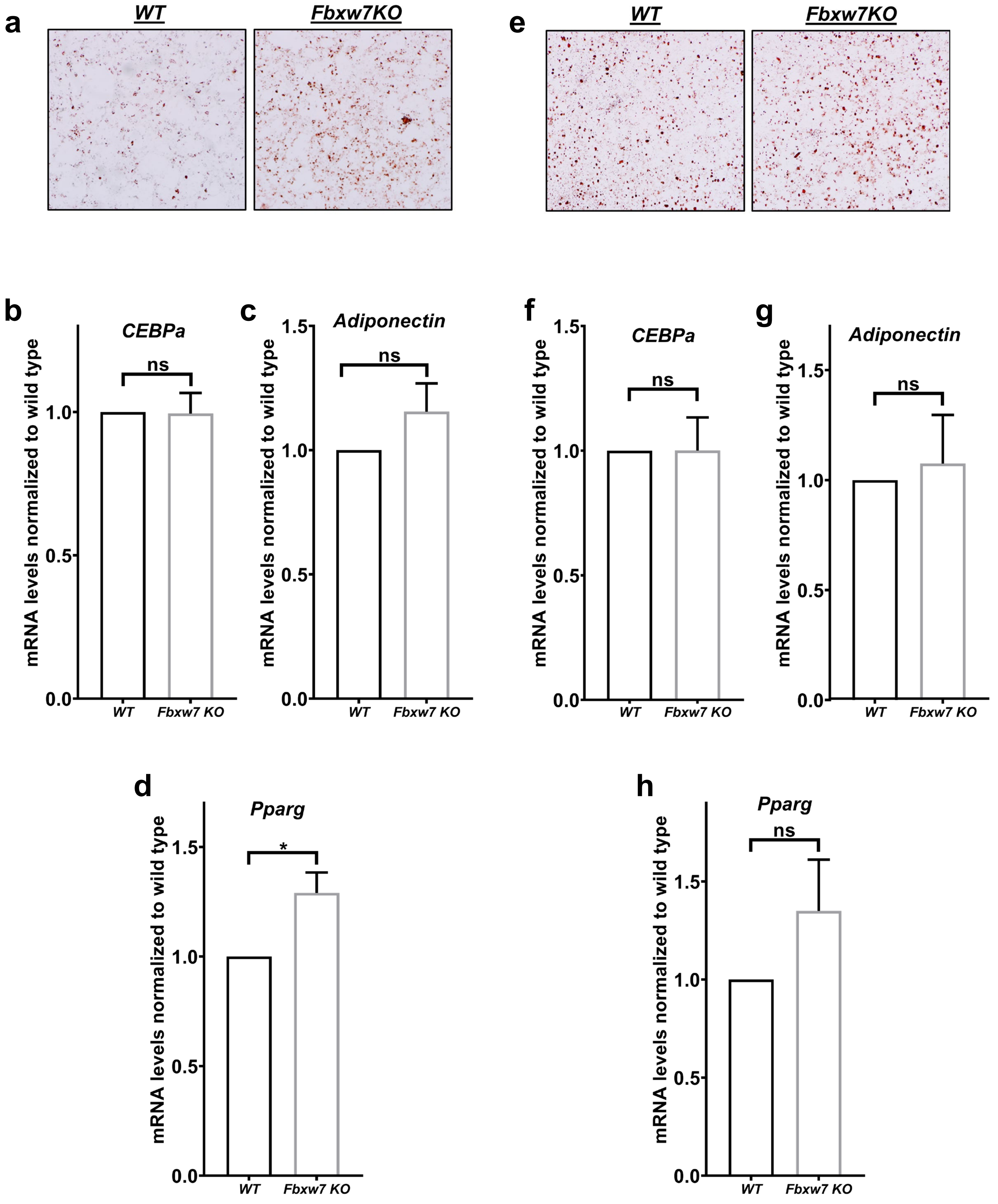

a
WT
Fbxw7KO
e
WT
Fbxw7KO
b
CEBPa
ns
mRNA levels normalized to wild type
1.0
0.5
0.0
WT
Fbxw7 KO
c
Adiponectin
ns
mRNA levels normalized to wild type
1.5
1.0
0.5
0.0
WT
Fbxw7 KO
f
CEBPa
ns
mRNA levels normalized to wild type
1.5
1.0
0.5
0.0
WT
Fbxw7 KO
g
Adiponectin
ns
mRNA levels normalized to wild type
1.5
1.0
0.5
0.0
WT
Fbxw7 KO
d
Pparg
*
mRNA levels normalized to wild type
1.5
1.0
0.5
0.0
WT
Fbxw7 KO
h
Pparg
ns
mRNA levels normalized to wild type
1.5
1.0
0.5
0.0
WT
Fbxw7 KO

Suppl. Fig 4

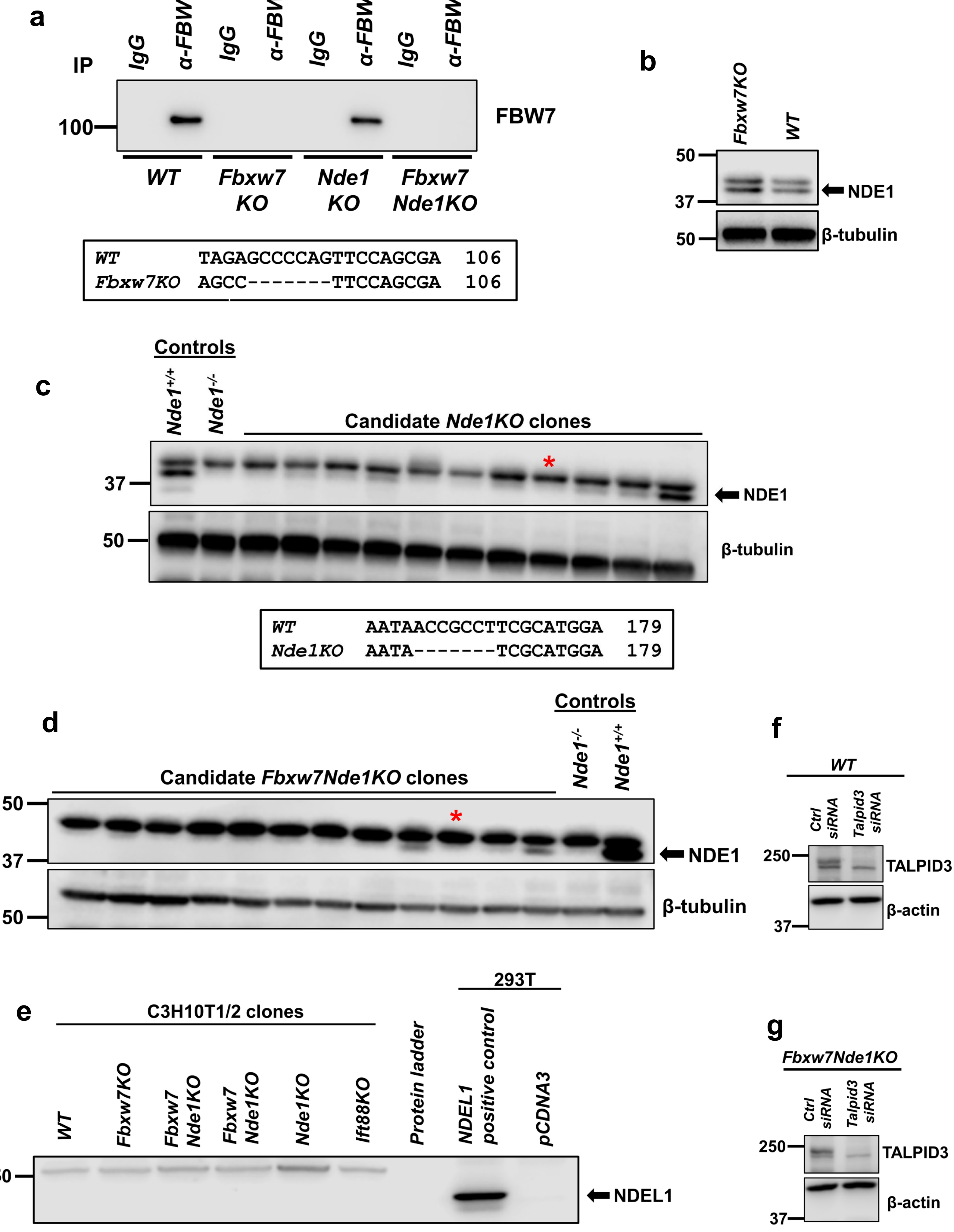
a
IP
IgG
α-FBW7
IgG
α-FBW7
IgG
α-FBW7
IgG
α-FBW7
100
FBW7
WT
Fbxw7 KO
Nde1 KO
Fbxw7 Nde1KO
WT TAGAGCCCCAGTTCCAGCGA 106
Fbxw7KO AGCC-------TTCCAGCGA 106
b
Fbxw7KO
WT
50
37
NDE1
50
β-tubulin
Controls
c
Nde1+/+
Nde1-/-
Candidate Nde1KO clones
*
37
NDE1
50
β-tubulin
WT AATAACCGCCTTCGCATGGA 179
Nde1KO AATA-------TCGCATGGA 179
Controls
d
Candidate Fbxw7Nde1KO clones
Nde1-/-
Nde1+/+
50
*
37
NDE1
β-tubulin
50
f
WT
Ctrl siRNA
Talpid3 siRNA
250
TALPID3
β-actin
37
e
C3H10T1/2 clones
293T
WT
Fbxw7KO
Fbxw7 Nde1KO
Fbxw7 Nde1KO
Nde1KO
Ift88KO
Protein ladder
NDEL1 positive control
pCDNA3
50
NDEL1
g
Fbxw7Nde1KO
Ctrl siRNA
Talpid3 siRNA
250
TALPID3
β-actin
37

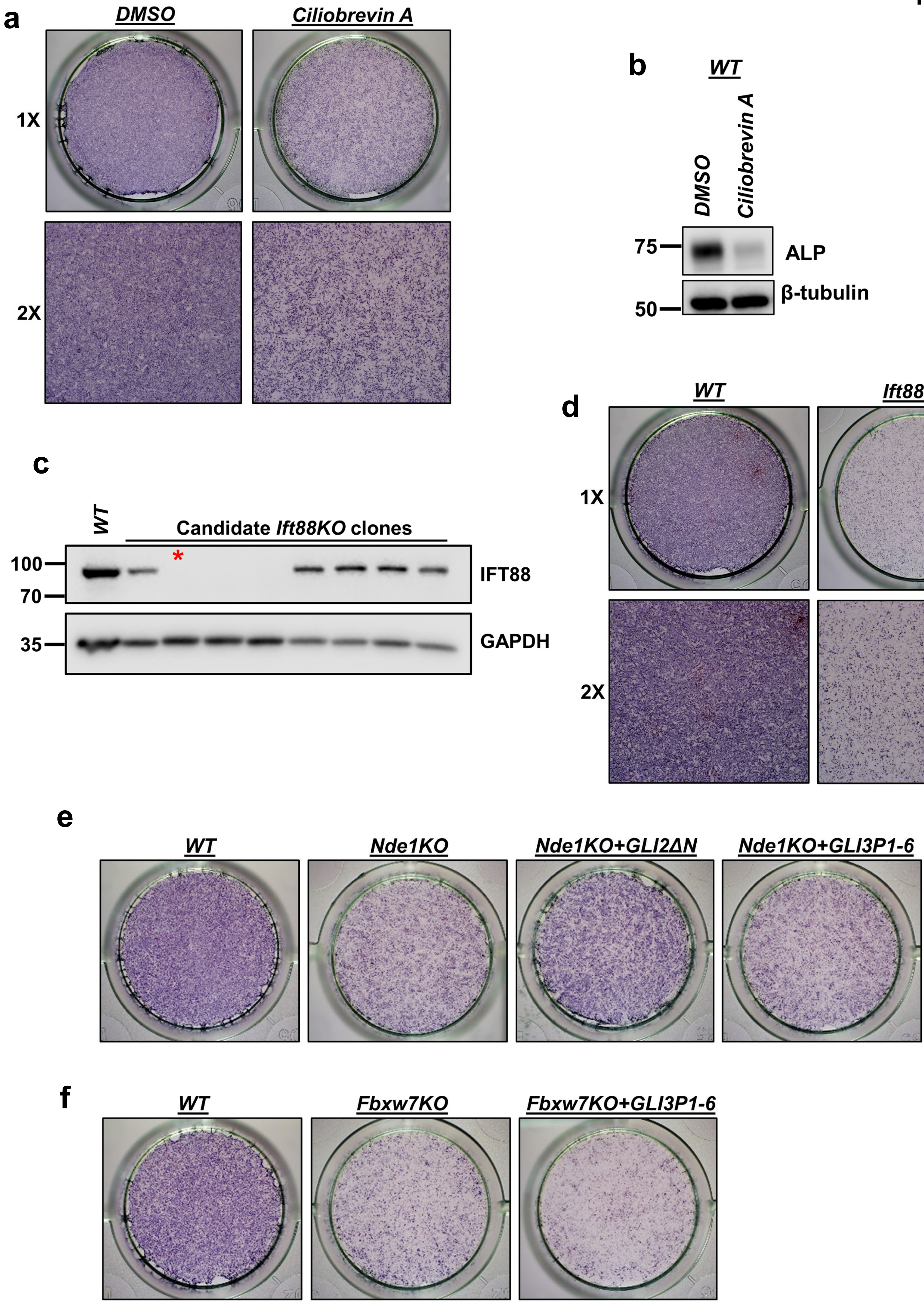
a
DMSO
Ciliobrevin A
1X
2X
b
WT
DMSO
Ciliobrevin A
75
ALP
β-tubulin
50
c
WT
Candidate Ift88KO clones
*
100
70
IFT88
35
GAPDH
d
WT
Ift88KO
1X
2X
e
WT
Nde1KO
Nde1KO+GLI2ΔN
Nde1KO+GLI3P1-6
f
WT
Fbxw7KO
Fbxw7KO+GLI3P1-6

**Suppl. Fig 6**

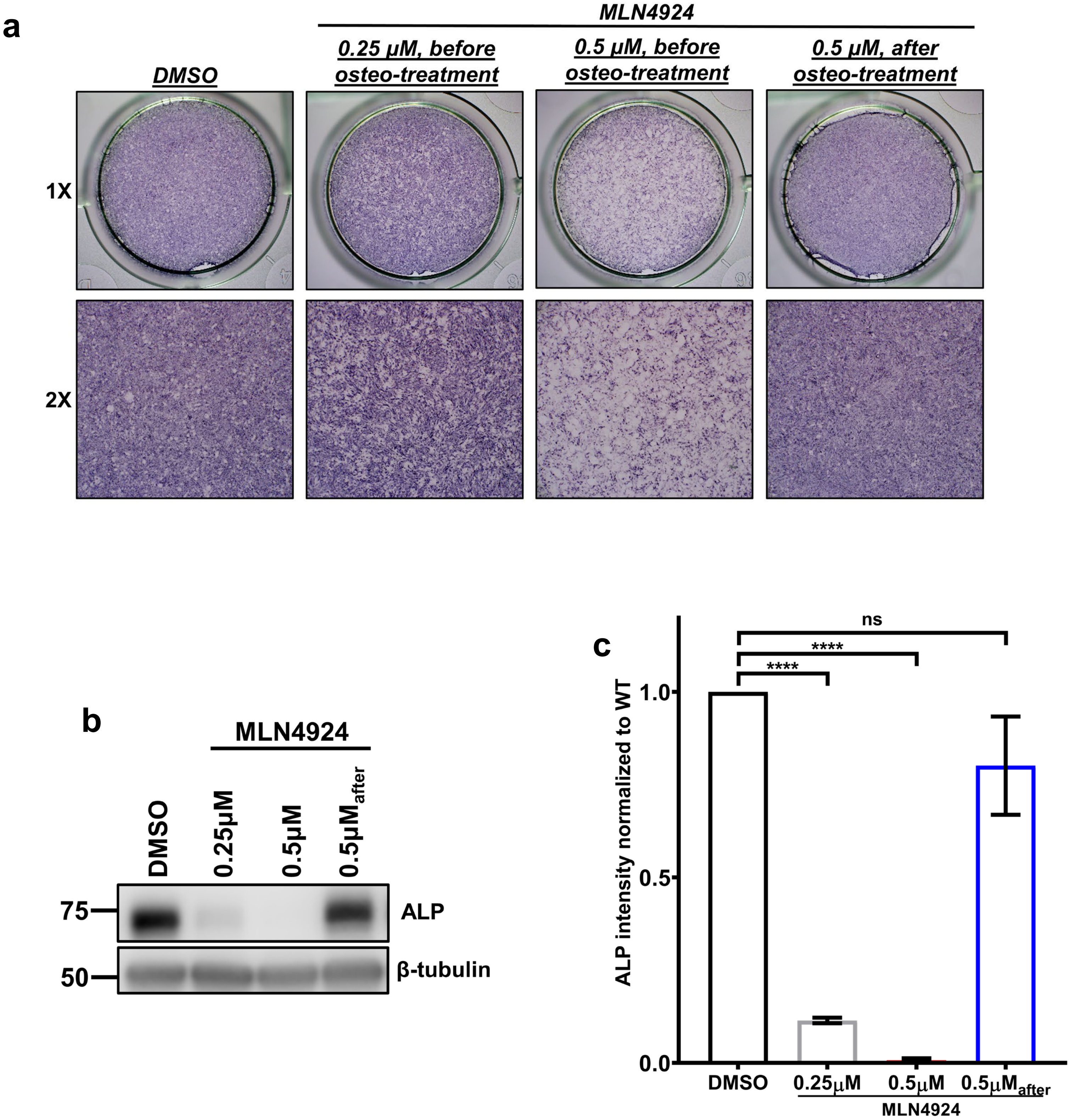

a
MLN4924
DMSO
0.25 µM, before osteo-treatment
0.5 µM, before osteo-treatment
0.5 µM, after osteo-treatment
1X
2X
b
MLN4924
DMSO
0.25µM
0.5µM
0.5µMafter
75
50
ALP
β-tubulin
c
ns
****
****
ALP intensity normalized to WT
1.0
0.5
0.0
DMSO
0.25µM
0.5µM
0.5µMafter
MLN4924
